# DUSP6 is transcriptionally upregulated by activated ALK and cooperates with ALK signaling to reduce lorlatinib sensitivity in neuroblastoma cells

**DOI:** 10.64898/2026.06.07.730727

**Authors:** Elliott Thompson, Vruti Patel, Christina Karapouliou, Vinothini Rajeeve, Pedro R. Cutillas, Andrew W. Stoker

## Abstract

Neuroblastoma is a pediatric, sympathoadrenal tumour accounting for 7-10% of childhood malignancies. Some neuroblastomas are driven by activating mutations in ALK kinase and inhibitors show promise in clinical trials. Nevertheless, with resistance an ever-present concern, it remains important to understand better the effectors and modulators of ALK signaling. Wild type ALK promotes ERK activation, raising expression of negative regulators such as the dual-specificity phosphatase (DUSP) DUSP6. DUSP6 though can be pro- or anti-oncogenic in different cancers and its role in neuroblastoma cells remains unclear. We sought to understand its role in cells with either wild type or mutated ALK. Mutated ALK strongly promotes *DUSP6* transcription, but apparently not through the ERK pathway. DUSP6 also appears to promote neuroblastoma cell proliferation without affecting ERK. Additionally, when DUSP6 is lost, the cells become more sensitive to ALK inhibitors lorlatinib and crizotinib. Phosphoproteomic analysis of such cells demonstrates that mutated ALK cooperates with DUSP6 to maximise signaling through several potential pathways, but again not through ERK or AKT. Their cooperation may also maintain optimal levels of N-Myc in MYCN-amplified neuroblastoma cells. While key substrates of DUSP6 remain to be determined in neuroblastoma cells, our study defines a novel role for this phosphatase in supporting the action of oncogenic ALK.

## 1. Introduction

Neuroblastoma is the most common extracranial tumour diagnosed in children [1]. The clinical outcomes for patients with neuroblastoma are diverse, where some tumours spontaneously regress without treatment whilst others present as an aggressive and progressive, metastatic disease. Patients with lower grade tumours display a 5-year survival greater than 90% [2], whereas high-risk patients still only have a 5-year survival of 40-50% even after multimodal treatments [3]. Hence there remains an urgent need to increase our understanding of oncogenic driver mechanisms in neuroblastoma and identify new therapeutic opportunities.

The most common oncogenic aberration in neuroblastoma is *MYCN* amplification, occurring in 20% of patients and associating with poor prognosis and metastasis [4]. Diverse therapies are being explored to perturb MYCN activity, with clinical trials ongoing [5]. The *ALK* oncogene is also mutated or amplified in 18% of high-risk neuroblastoma cases and is an independent predictor of overall survival [6]. Tumours where MYCN is amplified alongside mutations in ALK, have particularly poor prognosis. A range of ALK inhibitors have been generated that bind the ALK ATP-binding pocket [7] and clinical trials with these inhibitors show some promise in early clinical trials, albeit with complications due to resistance [8, 9]. A greater understanding of signalling components that cooperate with ALK could therefore reveal alternative combinatorial therapies that could be used alongside ALK inhibitors.

The ALK receptor can activate a range of signaling pathways, the most characterised being those centered on PI3K/AKT, STAT/JAK and RAS/ERK [9]. Long-term treatment of ALK-mutated neuroblastoma cells with ALK inhibitors leads to de novo mutations in NRAS and upregulation of ERK activity [9, 10]. Similar results are observed clinically where a greater frequency of ERK pathway mutations is found in relapsed neuroblastoma, highlighting the importance of ERK to ALK function in high-risk neuroblastoma [11]. As well as altering ERK activity, ALK-mutated neuroblastoma cells also increase the expression of negative feedback regulators of ERK, suggesting manipulation of ERK expression could alter ALK inhibitor therapies [12]. Although suppressing the negative regulators of ERK would appear counterintuitive, there are cases where cancer cells become addicted to negative feedback of ERK. In pre-B acute lymphoblastic leukaemia transformation, negative ERK regulators DUSP6, ETV5 and SPRY2 were key to robust oncogenic transformation [13]. Similarly, in NRAS and BRAF mutant melanoma cells there is a dual dependence on DUSP4 and DUSP6 activity to maintain appropriate but moderate levels of activated ERK, driving cell proliferation [14]. DUSP6 has been shown to have a pro-oncogenic role in several cancer types [15–19] and is recently shown to enhance cisplatin resistance in nasopharyngeal cancer cells [20] and can overcome HER2 inhibitor-induced dormancy in breast cancer cells [21]. ALK itself is also targeted by the phosphatases PTPN1 and PTPN2, and altering the expression of these phosphatases regulates ALK inhibitor efficacy [22]. Taken together, we hypothesised that if ERK-specific DUSP phosphatases were similarly pro-oncogenic in neuroblastoma cells, they may also influence the effectiveness of ALK inhibitors. In this study, we show that DUSP4, DUSP5 and DUSP6 are under the transcriptional control of ALK, and a detailed examination of DUSP6 shows that it can be regulated by ALK in an ERK-independent manner. We demonstrate that enforced loss of DUSP6 modestly suppresses cell growth in KELLY cells and these cells become more sensitive to ALK inhibition. Surprisingly, this DUSP6 loss does not appear to greatly impinge on the regulation of ERK signalling in these cells. This was confirmed through a phosphoproteomic screen, from which several further pathways of interest to neuroblastoma were shown to be influenced conjointly by DUSP6 loss and ALK inhibition. The study therefore suggests that DUSP6 may enhance ALK’s oncogenic potential by working cooperatively with ALK in ALK-mutated neuroblastoma cells.

## 2. Methods

### 2.1 Cell culture and chemicals

KELLY (CVCL_209) and SK-N-AS cells (CVCL_1700) were a gift from Prof. Frank Speleman, University of Ghent. IMR-32 cells (CVCL_0346) and SH-SY5Y (CVCL_0019) were supplied by ATCC, LAN-5 (CVCL_0389) were supplied by the Children’s Oncology Group Cell Culture and Xenograft Repository (Texas Tech University Health Sciences Center, USA), and LAN-1 cells (CVCL_1827) were a gift from Prof. John Anderson, UCL, London. All cell lines have been STR validated in the past year using the services of Northgene^TM^ (UK). Cells were incubated at 37 °C and 5% C0_2_ in a humidified incubator, in RPMI medium + GlutaMAX™ (ThermoFisher Scientific, Loughborough, UK) containing 10% FBS, 100 U/mL Penicillin, 0.1 mg/ml Streptomycin and 25 mM HEPES. Cells were routinely tested for mycoplasma using the MycoAlertTM PLUS Mycoplasma Detection Kit (Lonza, Slough, UK). Lorlatinib (PZ0039), Crizotinib (PZ0191), MG132 (M7449), EGF (SRP3027) and doxycycline (dox) (D1822) were purchased from Merck (Gillingham, UK). U0126 (SM106) was purchased from Cell Guidance Systems. Cyclohexamide was a gift from Prof. Owen Williams (University College London, UK). Chemical treatments were performed 24 hours after seeding cells.

### 2.2 Generation of DUSP6-null populations

Depletion of DUSP6 in KELLY and IMR-32 cells was achieved using an inducible CRISPR-Cas9 system described in Thompson et al. 2022 [23]. Briefly, inducible KELLY and IMR-32 populations (iKELLY and iIMR-32) were generated where Cas9 expression was under the control of a dox-inducible promoter. A *DUSP6* gRNA vector generated by cloning the gRNA sequence into plasmid pU6-sgRNA EF1Alpha-puro-T2A-BFP (Addgene) This was transfected into iKELLY and iIMR-32 and selected with puromycin. This resulted in Cas9-inducible clones stably expressing the gRNA (KELLY-G and IMR-32-G, respectively). KELLY-G and IMR-32-G were then treated with 0.5 μg/mL dox for 6 days to generate a mixed population of edited cells (KELLY-G+dox and IMR-32-G+dox). Following this, single cell cloning of mixed populations resulted in individual clones containing indels in the *DUSP6* gene, termed KELD6-1, KELD6-2 and KELD6-3. A flow chart is shown in Supplementary Figure 1.

### 2.3 Generation of DUSP6-overexpression population

Exogenous DUSP6 protein was re-introduced into KELLY and KELD6-2 cells using a GenEZ™ ORF Clone Vector (Genscript) containing the DUSP6 protein coding sequence. Cells were transfected with the *DUSP6* vector or an empty vector, pCI-Neo (Promega), using Lipofectamine 2000 (ThermoFisher Scientific) and selected for using G418 over 4 weeks. Overexpression of DUSP6 was verified via Western blot.

### 2.4 Immunoblotting

Whole cell lysates were prepared in lysis buffer (50 mM Tris Base; 150 mM NaCL; 1% Triton X-100; 0.02% Sodium Azide; 1 mM protease inhibitor cocktail (Merck Life Science UK Ltd); 1 mM sodium orthovanadate; 25 mM sodium fluoride). Protein concentration was determined using the Bradford Assay (Bio-Rad, Watford, UK). Samples were boiled and resolved by SDS-PAGE on 10% polyacrylamide gels. Protein was transferred to PVDF membranes and blocked in Tris Buffered Saline (15.4 mM Trizma-HCL; 4.62 mM Tris-base, 150 mM NaCl; pH 7.6) containing 10% (w/v) milk powder (Merck Life Science UK Ltd). Membranes were incubated overnight in primary antibody at 4°C and with secondary antibody conjugated to HRP for 1 hr (Agilent Technologies, Stockport, UK). Primary antibodies from Cell Signalling (Leiden, The Netherlands): phospho-ERK (pERK) (#9106; 1 : 1000); total ERK (tERK) (#9102; 1 : 2000); phospho-AKT (pAKT) (#4060; 1 : 1000); total AKT (tAKT) (#9272; 1 : 2000); pp38 (#9211; 1 : 1000); tp38 (#9212; 1 : 2000); MYCN (#9405; 1:1000), GAPDH (#2118; 1 : 10 000). DUSP6 antibody (ab76310) was from Abcam (Cambridge, UK). HRP was detected using ECL prime (Cytiva, Amersham, UK) and band intensities were quantified using Image Studio Lite (v5.2; LI-COR, Lincoln, NE, USA). Membranes were stripped in 0.2M NaOH for 20 minutes at 37 °C and then 20 min at room temperature before re-blocking membranes.

### 2.5. Phosphoproteomics

Phosphoproteomics experiments were performed using mass spectrometry as reported [24, 25]. In brief, iKELLY and KELLY-G cells were treated with 0.5 µg/ml dox for 6 days and then without dox for 7 – 9 days. Cells were treated with 30 nM lorlatinib for 1 hr before lysis in: 8 M urea; 1 mM sodium orthovanadate; 1 mM sodium fluoride; 1 mM β-glycerol phosphate; 2.5 mM Disodium diphosphate. Proteins were digested into peptides using trypsin as previously described [26, 27]. Phosphopeptides were enriched from total peptides by TiO2 chromatography essentially as reported previously [28]. Dried phosphopeptides were dissolved in 0.1% TFA and analysed by nanoflow ultimate 3000 RSL nano instrument coupled online to a Q-Exactive plus mass spectrometer (ThermoFisher Scientific). Gradient elution was from 3% to 28% solvent B in 90 min at a flow rate of 250 nL·min^−1^ with solvent A being used to balance the mobile phase (solvent A was 0.1% formic acid in water and B was 0.1% formic acid in acetonitrile). The spray voltage was 1.95 kV and the capillary temperature was set to 255 °C. The Q-Exactive plus was operated in data-dependent mode with One Survey MS scan followed by 15 MS/MS scans. The full scans were acquired in the mass analyser at 375–1500 m/z with the resolution of 70 000, and the MS/MS scans were obtained with a resolution of 17 500.

MS raw files were converted into Mascot Generic Format using mascot distiller (version 2.6.1; Matrix Science, London, UK) and searched against the SwissProt (release Sep 2018) restricted to human entries using the mascot search daemon (version 2.6.0; Matrix Science). Allowed mass windows were 10 ppm and 25 mmu for parent and fragment mass to charge values, respectively. Variable modifications included in searches were oxidation of methionine, pyro-glu (N-term) and phosphorylation of serine, threonine and tyrosine. All raw data are available at 10.17632/f2jx8t4x8h.1 (Mendeley Data).

Kinase substrate enrichment analysis (KSEA) was performed on each dataset as described before by calculating a z-score between the distribution of changes in phosphorylation of substrates for given kinases relative to the distribution of all fold changes [24].

### 2.6 Cell growth and viability assays

Resazurin (sodium salt; Merck Life Sciences, UK) was used to measure cell growth and viability. Briefly, neuroblastoma cells were seeded at 2 x 10^3^ cells per well in 96-well plates. Chemical treatments were performed after 24 hours and cells were incubated for 6 days. Resazurin was dissolved in PBS and added to cells at a concentration of 0.0015% (w:v). Cells were incubated for 4 hours and fluorescence intensity was measured on a FLUOstar Optima (BMG Labtec Ltd., Aylesbury, UK).

### 2.7 Quantitative PCR

RNA was extracted from neuroblastoma cells using the mini RNeasy kit (Quigen). RNA was reverse-transcribed to cDNA using the Transcriptor High Fidelity cDNA Synthesis Kit (Roche). CDNA was mixed with 10 µM forward primer, 10 µM reverse primer and the iTaq Universal SYBR Green Supermix (Bio-Rad, Watford, UK) and qRT-PCR reactions were run on a CFX Connect PCR machine (Bio-Rad, Watford, UK). Primers used are presented in Table 1.

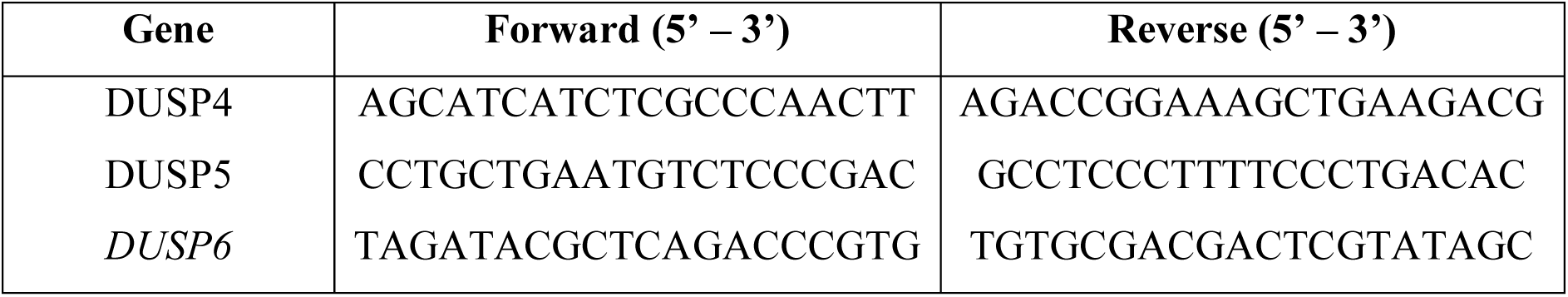

### 2.8 Statistical analysis

To assess the statistical significance between groups an independent t-test was performed. Where multiple groups were compared, a one-way ANOVA was performed with a Dunnett post-hoc analysis. Statistical analysis was performed on SPSS (v28; IBM, Portsmouth, UK).

## 3 Results

### 3.1 DUSP6 influences cell growth without altering MAP kinase activation

Cell growth- and survival-promoting roles for DUSP6 have been identified in a range cancers [15–19, 29]. Our initial aim was therefore to determine if DUSP6 supported or suppressed neuroblastoma cell growth in culture. We have previously generated neuroblastoma cell lines with dox-inducible Cas9 (iKELLY and iIMR32; [23]). Stable transfection of these with gRNAs targeting *DUSP6* produced mixed populations KELLY-G and IMR32-G. When KELLY-G and IMR32-G were treated with dox, this generated mixed populations with a range of indels in *DUSP6. DUSP6*-null subclones of these are named KELD6-1, KELD6-2 and IMRD6-1 [23] with undetectable level of DUSP6 (see Fig 8 in [23] and flow chart in Supplementary Figure 1). To determine if loss of DUSP6 altered cell proliferation, the iKELLY and iIMR32 cells were treated with dox alongside the KELLY-G and IMR32-G cells (Figure 1A). Although modest, there was a statistically significant reduction in cell growth in the KELLY-G+dox population compared to the iKELLY+dox population (Figure 1A). The IMR32-G-dox population growth did not differ from iIMR32-dox cells. The growth of two *DUSP6*-null subclones KELD6-1 and KELD6-2 were compared to the parental KELLY-G population. There was a significant decrease in one but not the other subclone. These data suggest that DUSP6 can have growth-promoting effects, but this is variable both between cell lines KELLY and IMR32, and between subclones with indels (Figure 1B).

**Figure 1.**
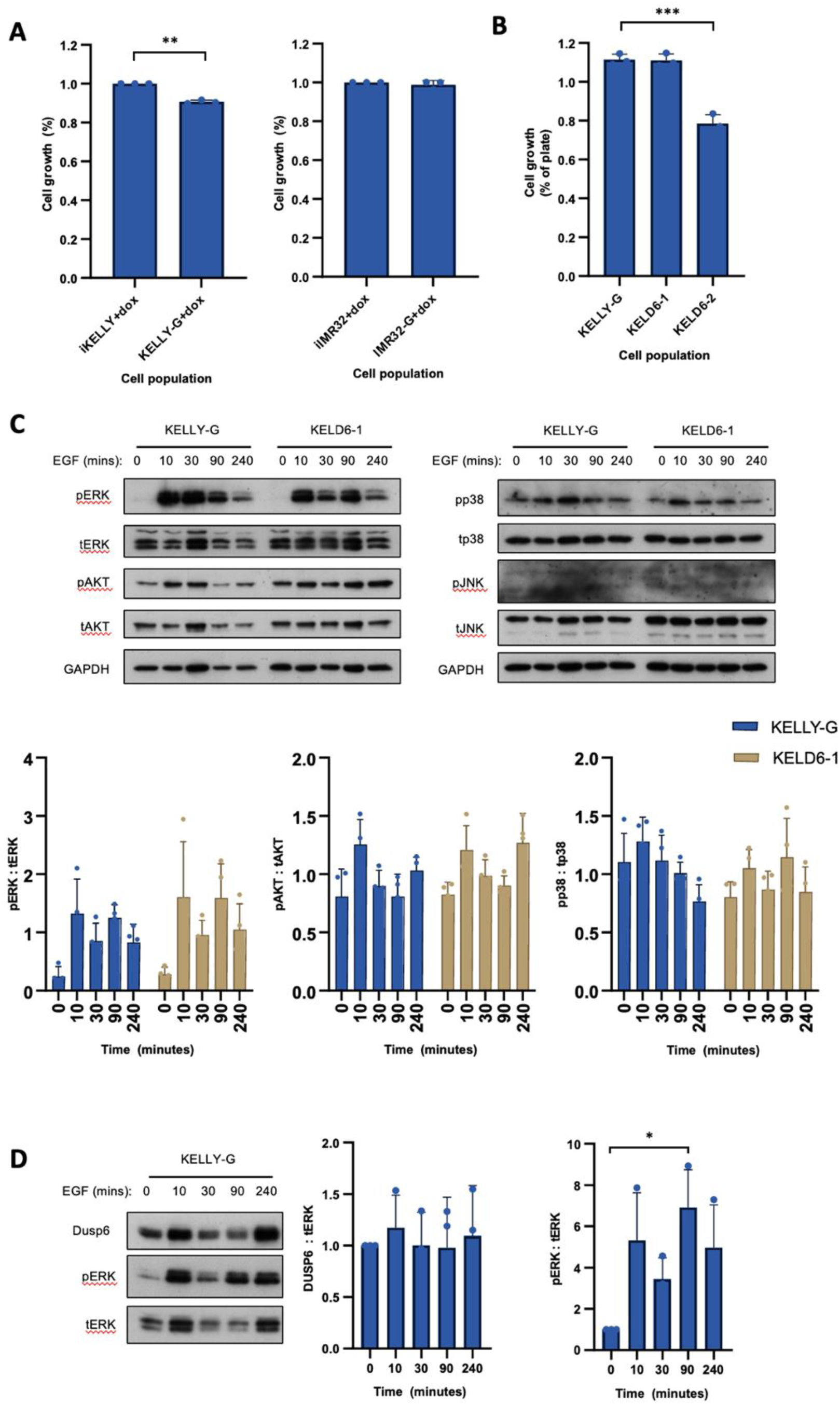
Loss of DUSP6 can reduce neuroblastoma cell growth but does not affect ERK activity. (A) Cell growth of mixed populations of KELLY and IMR-32 cells without *DUSP6* gRNA (iKELLY and iIMR32) and with *DUSP6* gRNA (KELLY-G and IMR32-G), after treatment with 0.5 µg/ml dox for 6 days; see [23] for DUSP6 knockdown data. (B) Cell growth of KELLY-G cells, not treated with dox, compared to *DUSP6*-KO subclones (KELD6-1 and KELD6-2; [23]. Cell growth was measured after 6 days. Data are expressed as a mean ± SD (n≥3). One-way ANOVA compared to (A) iKELLY or (B) KELLY-G; ***p ≤ 0.0005. (C) KELLY-G cells (not treated with dox), and KELD6-1 cells were treated with 20 ng/ml EGF for 10, 30, 90 and 240 minutes and then assessed for phosphorylated ERK, JNK, p38 and AKT expression (n≥3). DUSP6 expression was normalised to GAPDH. One-way ANOVA compared equivalent timepoints in *DUSP6* WT and KO cells. (D) KELLY-G cells (not treated with dox), were treated with 20 ng/ml EGF for 10, 30, 90 and 240 minutes and then assessed for DUSP6 levels. DUSP6 expression was quantified and normalised to total ERK. One-way ANOVA compared to untreated cells; *p ≤ 0.05; n=3.

Considering DUSP6 is normally deemed a key ERK-phosphatase, we examined if DUSP6 could influence the dynamics of ERK signalling in KELLY cells. KELLY-G cells (*wtDUSP6*) and *DUSP6*-null clone KELD6.1 were treated with EGF and these were assessed for ERK, p38, JNK and AKT activity (Figure 1C). Strong increases in pERK were observed with EGF, while pAKT and pp38 were not consistently elevated. Surprisingly, the response of all three proteins to EGF treatment was comparable in KELLY-G and *DUSP6*-null cells, suggesting DUSP6 loss was not significantly altering ERK, p38 or AKT activity in KELLY cells. Although no clear changes were observed in any replicate for pJNK, these data should be treated with caution since the pJNK signals were reproducibly weak. Similar results were observed in a preliminary experiment using IMR32 cells (Supplementary Figure 2). A similar lack of change in pERK, pAKT or pp38 was seen in KELLY cells where DUSP6 protein was suppressed using siRNAs (data not shown).

In non-neuroblastoma cells, DUSP6 expression is documented to rise in response to ERK stimulation as part of a negative feedback loop [30, 31]. Parental and KELD6.1 cells were treated with EGF and DUSP6 was quantified (Figure 1D). ERK phosphorylation is increased by EGF (Figure 1D). However, DUSP6 protein expression, when normalised to total ERK levels, did not significantly change over time. Collectively, these data suggest that although DUSP6 may marginally promote KELLY cell growth, DUSP6 perturbation alone does not obviously influence MAPK signalling pathways in these cells. The well-defined DUSP6-ERK transcriptional feedback loop appears to be weak or may be compensated by other DUSP enzymes in these cells. This may also be true for IMR-32 cells which express wild type ALK (Supplementary Figure 2).

### 3.2 *DUSP6* expression is driven transcriptionally by ALK in an ERK-dependent and -independent manner

Considering the modest proliferation effect of DUSP6 loss on KELLY, alongside no effects in IMR-32, we hypothesised that the regulatory interactions between ALK and DUSP6 may be distinct in the two cell types due to their differing ALK mutation status (KELLY^F1174L^ and IMR-32^WT^). This was also suggested by a previous study that identified *DUSP4* as an ALK target gene [32]. To first evaluate whether DUSP family members are transcriptionally regulated by ALK, DUSP gene expression was extracted from two previously published bulk RNA sequencing datasets after treatment with the ALK inhibitor lorlatinib [32, 33] (Figure 2A). *DUSP4*, *DUSP5* and *DUSP6* were downregulated strongly in two cell lines, more weakly in another, but not significantly affected in ALK-WT cells, one of which was IMR-32. Similar transcriptional effects were found in CLB-GA cells using several ALK inhibitors [12]. We observed similar results in LAN5 and KELLY cells (Figure 2B), with *DUSP6* being the most affected by inhibition of ALK. This indicates that elevated expression of these genes is significantly dependent on aberrantly activated ALK. It was also noted that genes such as *DUSP2*, *DUSP9* and *DUSP10* were modestly upregulated in some but not all cell lines (Figure 2A), indicating that these genes are suppressed when ALK is activated in these tumour-derived cells.

**Figure 2.**
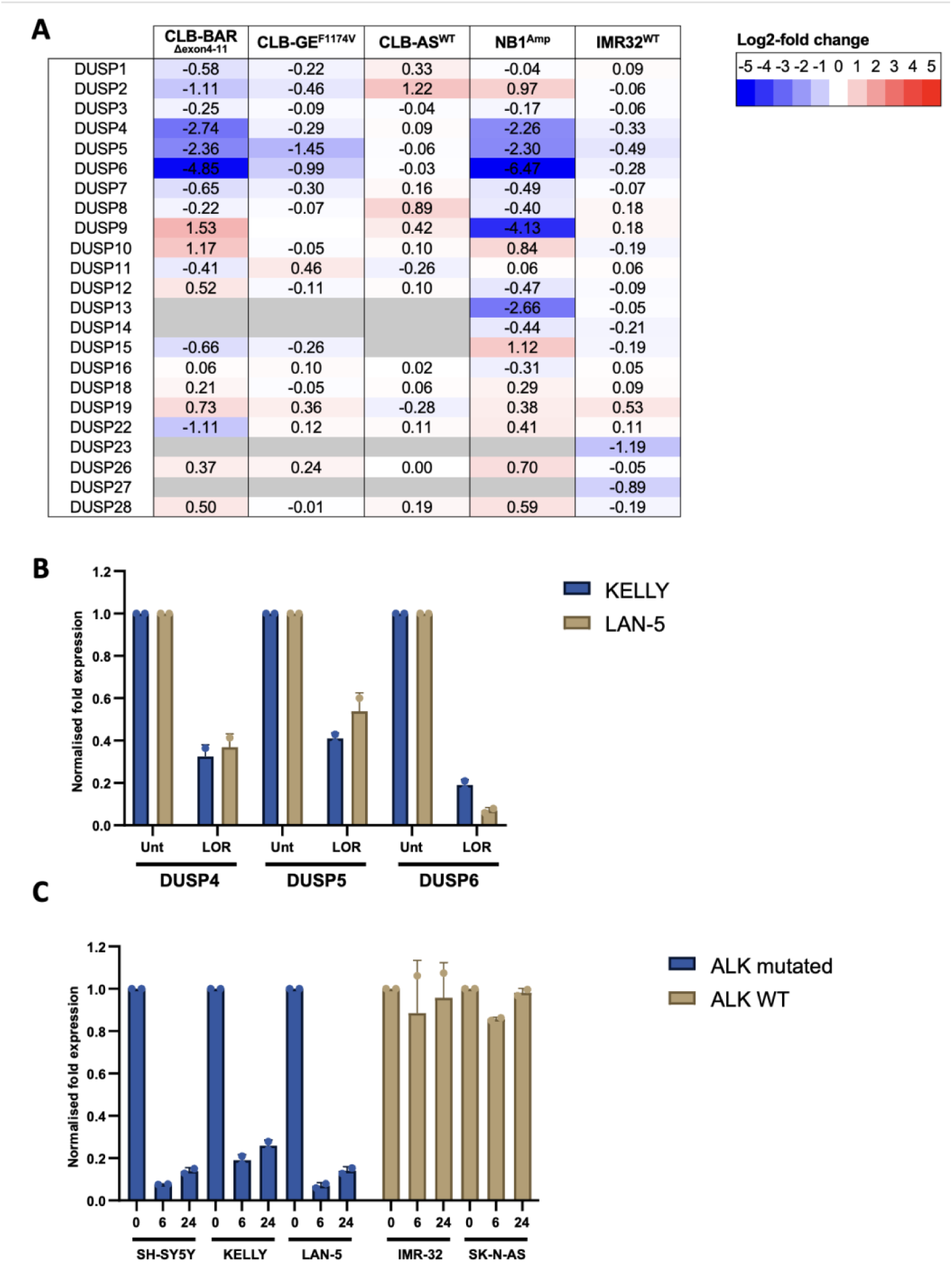
A range of DUSP family members are transcriptionally regulated by ALK. **A**, Log2-fold change in mRNA expression of DUSP family members after 24 hour treatment with 30 nM lorlatinib (LOR) in 5 neuroblastoma cell lines. CLB-BAR, CLB-GE and CLB-AS RNA sequencing data obtained from Van den Eynden et al., 2018; NB1 and IMR-32 RNA sequencing data were obtained from Borenas et al., 2021. ALK status denoted in superscript. **B**, KELLY and LAN-5 cells were treated with 30 nM LOR for 24 hrs and the expression of *DUSP4*, *DUSP5* and *DUSP6* was assessed by qPCR (n = 2). **C,** SH-SY5Y, KELLY, LAN-5, IMR-32 and SK-N-AS cells were treated 30 nM LOR for 6 and 24 hrs and *DUSP6* expression was assessed by qPCR (n =2). In B and C, data is normalised to the non-lorlatinib treated samples.

To further assess the dependence of DUSP6 expression downstream of ALK, an expanded set of ALK WT and mutated cell lines were treated with lorlatinib and assessed for *DUSP6* gene expression (Figure 2C). Within 6 hrs Lorlatinib treatment significantly inhibited *DUSP6* gene expression (>80%) in ALK mutated cells, supporting the hypothesis that ALK activity is acutely required for maintaining *DUSP6* expression. In wild type ALK-expressing cells, *DUSP6* expression is minimally affected by lorlatinib (Figure 2C).

We hypothesised that in ALK-activated cells, ALK may stimulate *DUSP6* expression via MEK-ERK signalling, since DUSP6 is a known, negative regulator of ERK [30, 31]. To test this, each cell population was treated with a MEK or ALK inhibitor and DUSP6 protein expression was assessed. MEK inhibition in KELLY cells does not suppress *DUSP6* mRNA to the same extent as ALK inhibition (Figure 3A), suggesting alternative routes for *DUSP6* gene regulation. This also appears to be the case at the protein level for ALK-mutated KELLY and LAN5, where MEK inhibition and ALK inhibition similarly suppress pERK (Figure 3B). In these cells though, MEK inhibition only reduces DUSP6 expression by 50%, whereas ALK inhibition further drives down DUSP6 levels >90%, suggesting that non-MEK/ERK signals from ALK can sustain DUSP6. In SH-SY5Y, ALK inhibition strongly blocks p-ERK, whereas MEK inhibition is only partial. DUSP6 however is not significantly suppressed by MEK inhibition, whereas it is greatly reduced after ALK inhibition. In SY5Y it also appears that ERK phosphorylation is at least partially maintained even when MEK is inhibited, and this is ALK-dependent. In IMR-32 and SK-N-AS, neither of which are ALK-mutated or addicted, the ALK inhibitor has no clear effect on pERK or DUSP6 levels, indicating that the wild type ALK has minimal influence over these proteins under these conditions. In IMR-32, ALK may even somewhat suppress DUSP6 given that DUSP6 level increase at least temporarily after LOR treatment (Figure 3B). MEK inhibition, however, does effectively suppress p-ERK and DUSP6 levels together. So in these non-ALK addicted cells, DUSP6 expression may be under more direct ERK control. Collectively these five cell lines have distinct circuits in place downstream of ALK to control DUSP6 levels (Figure 3C), reflecting the genetic and biochemical heterogeneity of neuroblastoma and neuroblastoma-derived cell lines. Of particular interest are the ALK-addicted lines, where DUSP6 expression is only partially dependent on MEK signals, but keenly maintained by the mutated ALK signaling.

**Figure 3.**
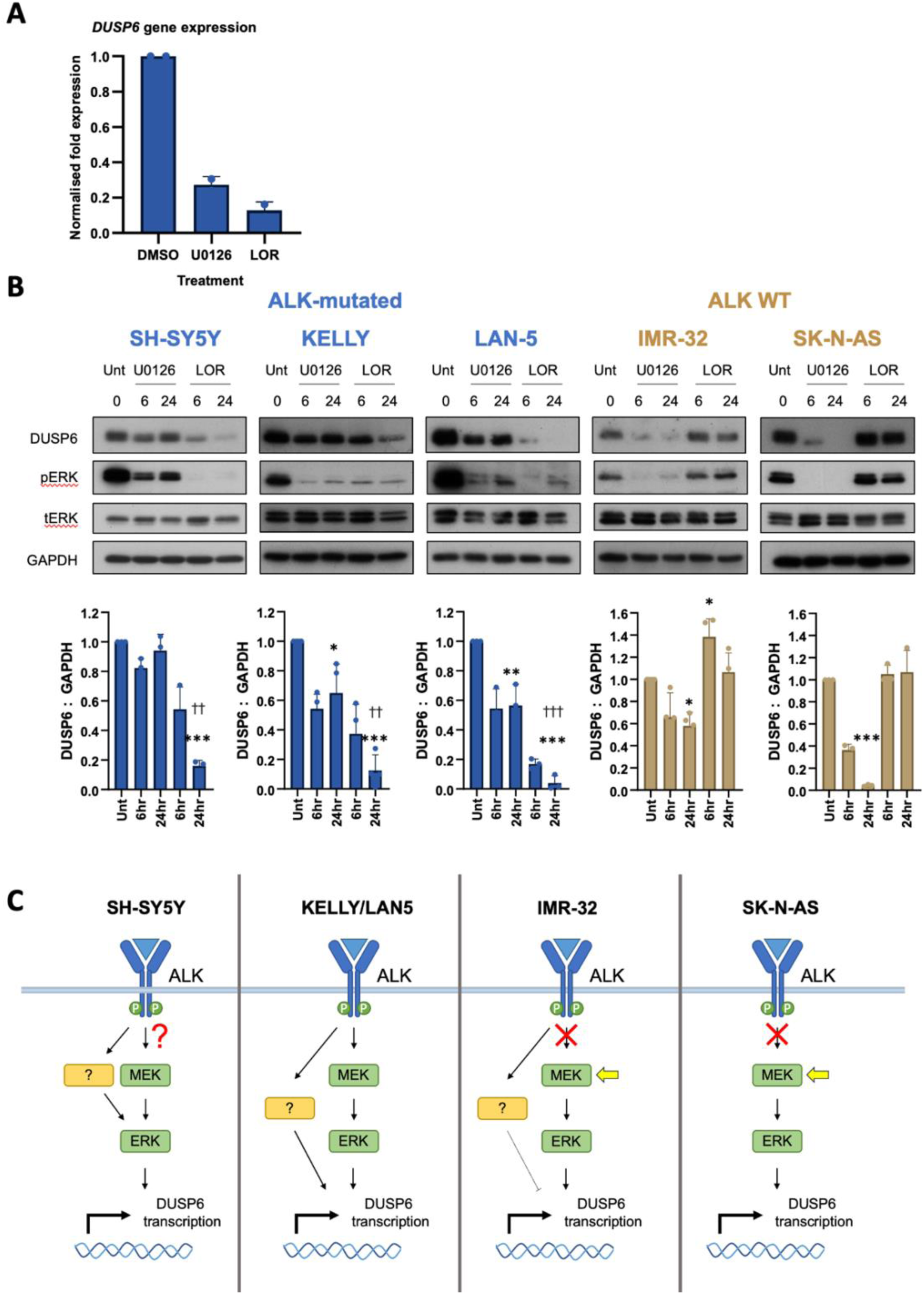
DUSP6 is regulated by ALK in an ERK-dependent and independent manner. (A) KELLY cells were treated with 30 nM LOR and 10 µM U0126 for 6 hours and *DUSP6* expression was assessed by qRT-PCR (n = 2). SH-SY5Y, KELLY, LAN-5, IMR-32 and SK-N-AS cells were treated with 30 nM LOR and 10 µM U0126 and assessed for DUSP6 expression and pERK levels via Western blot. DUSP6 and pERK were normalised to GAPDH and tERK respectively. One-way ANOVA comparing 24 hr timepoints to Untreated (Unt); *p ≤ 0.05, **p ≤ 0.005, ***p ≤ 0.0005. One-way ANOVA comparing 24 hr LOR timepoints to 24 hr U0126; ††p ≤ 0.005, †††p ≤ 0.0005. Triplicate experiments were performed. (C) Schematic of proposed models for ALK-mediated activation of DUSP6 expression in cell lines assessed. Red crosses and question marks indicate pathway links that are not detectable, or not yet clear, respectively. Yellow arrows indicate presence of alternative sources of MEK stimulation in these cells revealed under LOR treatment.

### 3.3 Post-translation control of DUSP6 protein levels

It is well documented that DUSP6 can be regulated post-translationally [34–36]. To investigate this here, cyclohexamide (CHX) chase assays were performed to assess the rate of DUSP6 turnover with or without lorlatinib (Figure 4A). DUSP6 turnover varied greatly between cell lines. In SK-N-AS, DUSP6 is degraded within 4 hours, demonstrating high protein instability, whereas in LAN-5 and IMR-32 cells the DUSP6 half-life was ∼4-6 hours. Conversely, in KELLY cells DUSP6 appeared more stable and required a 10-hour treatment for levels to reduce greater than 50% (Supplementary Figure 3). The addition of lorlatinib did not affect the rate of DUSP6 turnover in KELLY, IMR-32 or SK-N-AS. In LAN-5 cells it has a more complex effect, since ALK appears to suppress DUSP6 levels, but also promotes DUSP6 proteolysis since the addition of lorlatinib reduced DUSP6 turnover in the presence of CHX.

**Figure 4.**
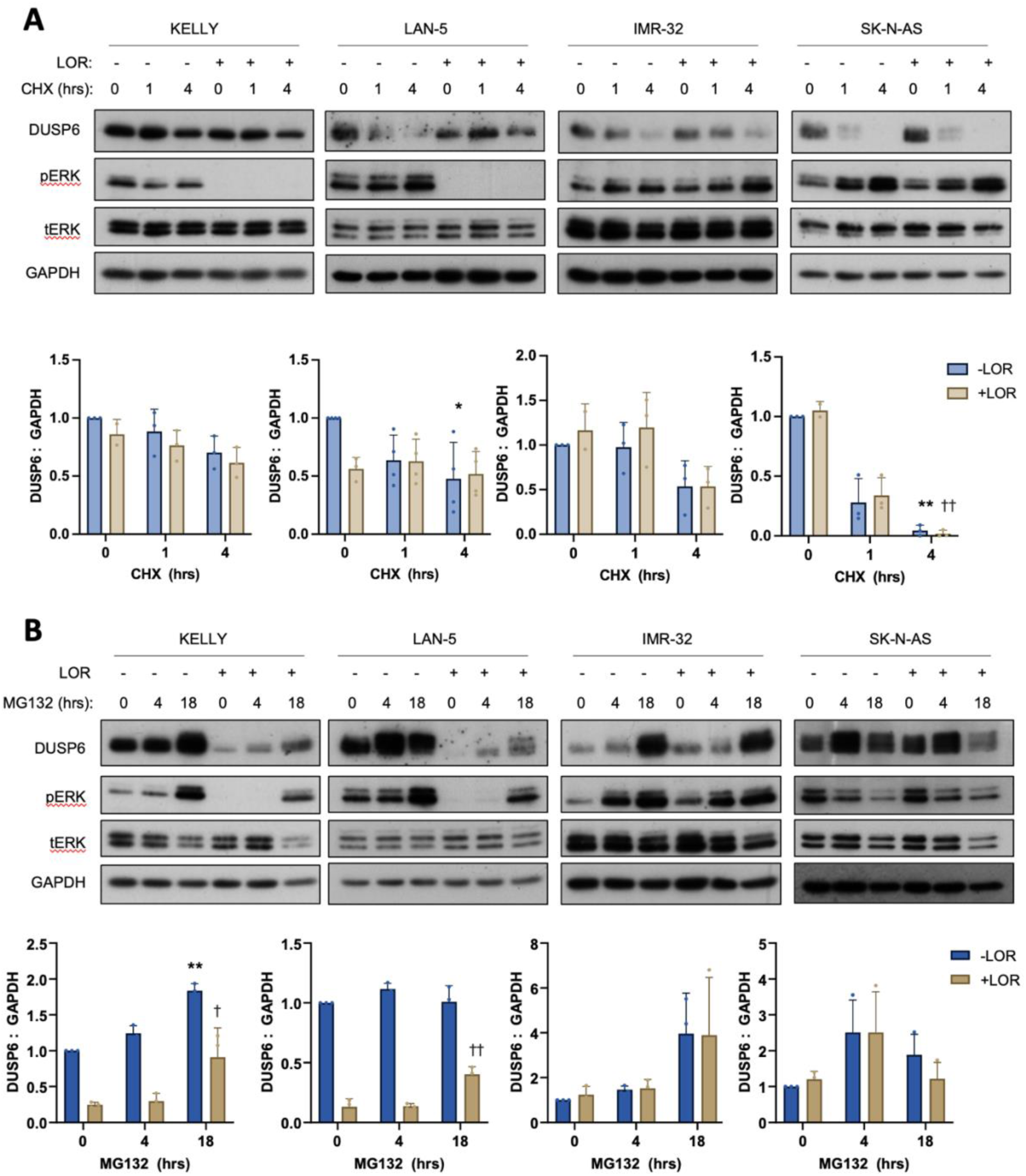
Post-transcriptional control of DUSP6. Neuroblastoma cell lines were treated with 40 µg/ml CHX for (A) 1 and 4 hours in the presence or absence of 30 nM lorlatinib and DUSP6 expression was assessed by Western blot (n≥3). (B) Neuroblastoma cell lines were treated with 10 mM MG132 for 4 and 18 hours in the presence or absence of 30 nM lorlatinib followed by immunoblotting for DUSP6 expression and pERK levels (n≥3). Lorlatinib treatment was kept constant at 18 hours. One-way ANOVA compared to Untreated (Unt); *p ≤ 0.05, **p ≤ 0.005. One-way ANOVA compared to 30nM lorlatinib; ^†^p ≤ 0.05, ^††^p ≤ 0.005. Triplicate experiments were performed.

We next assessed how the proteosome may regulate DUSP6 degradation. neuroblastoma cells were treated with the proteosome inhibitor MG132 with or without lorlatinib. In KELLY and IMR-32, treatment with MG132 led to an accumulation of DUSP6, suggesting DUSP6 is degraded via the proteosome (Figure 4B). In KELLY cells, although concurrent treatment with lorlatinib significantly reduced DUSP6 protein, DUSP6 still accumulated over time, suggesting DUSP6 proteolysis is not modulated by ALK signals directly. In LAN-5 cells, proteosome inhibition did not lead to accumulation of DUSP6 over 18 hours, suggesting DUSP6 is either not rapidly degraded via the proteosome under basal conditions, or ALK is able to maintain strong proteolysis at this level of MG132. The latter is supported by the fact that ALK inhibition in LAN5 appears to slow DUSP6 degradation during CHX treatment (Figure 4A), and double treatment with MG132 and ALK inhibitor allows DUSP6 levels to partially recover (Figure 4B). Taken together, these data suggest ALK does not regulate DUSP6 post-translationally, but instead operates predominantly at a transcriptional level. The exception is the possible promotion of proteolysis in LAN5 cells, although this is greatly counterbalanced by the transcriptional stimulation. It is also observed that ERK is activated by proteosome inhibition in KELLY, LAN5 and IMR-32, with or without ALK inhibition. ERK is however suppressed in SK-N-AS cells. There is some correlation between pERK and DUSP6 levels in KELLY and LAN5 when ALK is inhibited, possibly revealing some weak, positive control of DUSP6 by ERK in the absence of mutated ALK activity.

### 3.4 Complete loss of DUSP6 sensitises Neuroblastoma cells to ALK inhibition

If ALK is regulating and sustaining DUSP6 levels, it is conceivable that in turn DUSP6 is cooperating somehow with ALK signalling. We therefore assessed whether loss of DUSP6 affected KELLY cell growth in response to ALK inhibition. Dose-response survival assays were initially performed with lorlatinib and crizotinib in iKELLY cells (KELLY cells expressing dox-inducible Cas9; [23], and a mixed population KELLY-G cells (iKELLY cells expressing DUSP6-targeted guide RNA; [23] with depletion of DUSP6 (KELLY-G cells, Figure 8B in [23]. The gRNA-treated population showed approximately 20% reduction in IC50 to both drugs, but this did not reach statistical significance (Supplementary Figure 4). The same question was then addressed in *DUSP6*-null subclones (Figure 5A). Loss of DUSP6 in the KELD6-1 and KELD6-2 subclones significantly sensitised cells to ALK inhibition. Loss of DUSP6 more weakly sensitised the KELD6-3 subclone to lorlatinib, but not to crizotinib. The mixed population analysis (Supplementary Figure 4) is therefore likely to represent the average over a range of subclone responsiveness. A similar inhibitor analysis of IMR32-G and the DUSP6-depleted subclone IMRD6-1 showed no significant sensitising effect of DUSP6 depletion on inhibitor effectiveness (Supplementary Figure 5). To try to determine whether the sensitisation effect in KELLY was due to loss of DUSP6, the KELD6-2 population was transfected with a *DUSP6* overexpression vector. Validation of DUSP6 overexpression is demonstrated in Figure 5B where DUSP6 expression increased 3-fold above normal levels. Treatment with lorlatinib did not reduce exogenous DUSP6 expression, further indicating that the endogenous *DUSP6* gene is under tight transcriptional regulation by ALK in KELLY cells and that ALK signals do not control DUSP6 protein level post-translationally. Dose-response assays were performed with crizotinib and lorlatinib in the cells overexpressing DUSP6 (Figure 5C). There was an apparent modest reduction in the sensitisation to crizotinib, and small reduction with lorlatinib, although these did not reach statistical significance. Therefore the ability to rescue the sensitisation to inhibitors using overexpression of DUSP6 above endogenous levels is apparent but at best small.

**Figure 5.**
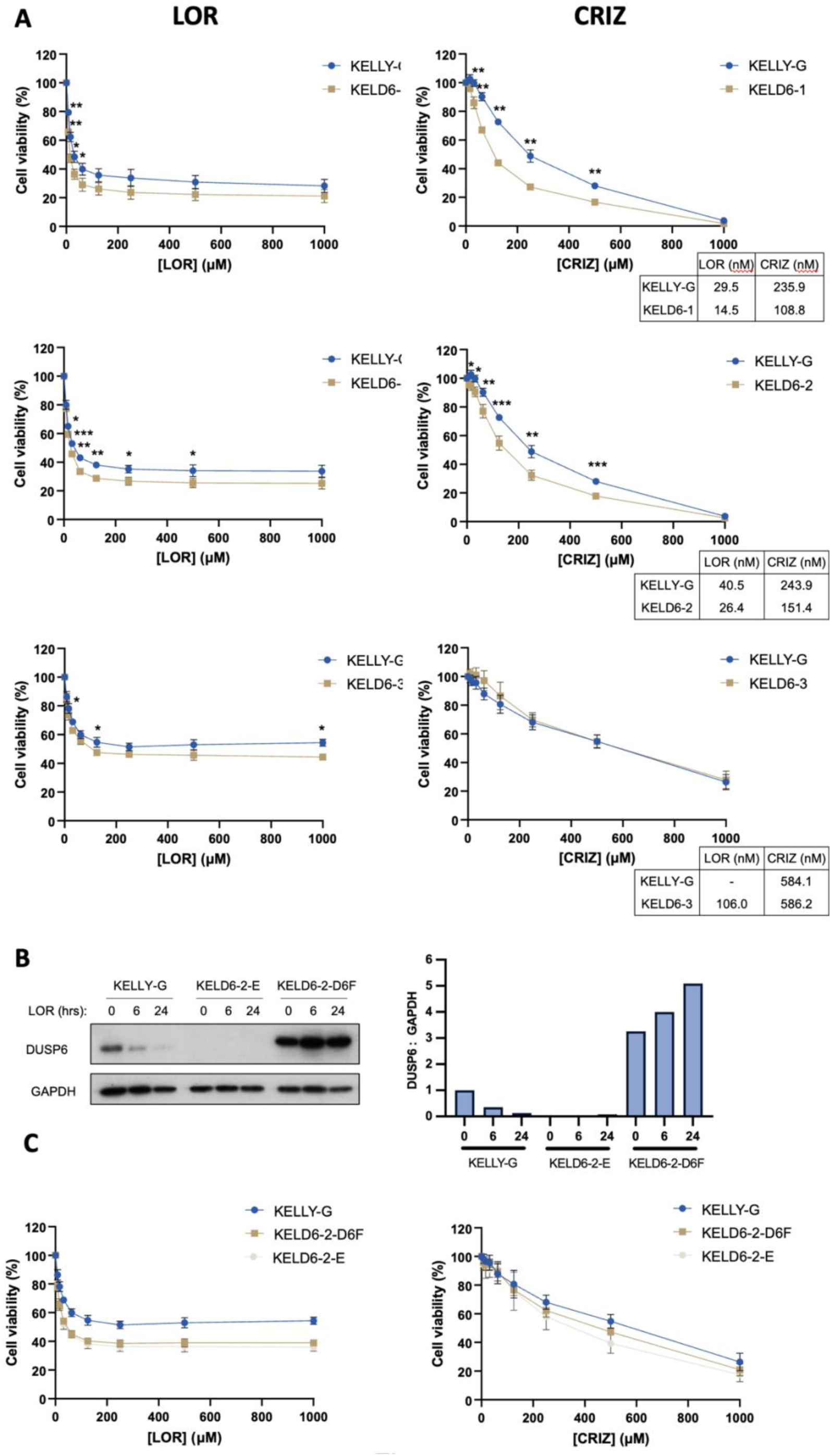
Loss of DUSP6 sensitises neuroblastoma cells to ALK inhibition. (A) KELLY-G and *DUSP6*-KO subclonal derivatives were treated with increasing concentrations of lorlatinib (LOR) and crizotinib (CRIZ) and a cell viability assay was performed. Data were normalised to cells treated with DMSO and then expressed as a mean ± SD (n≥3). Mean EC50 values are displayed alongside dose-response curves. An independent samples t-test was performed comparing equivalent drug concentrations; *p ≤ 0.05, **p ≤ 0.005, ***p ≤ 0.0005. (B) Western blot assessing DUSP6 expression in KELLY-G cells and KELLY cells treated with an empty (KELD6-2-E) and DUSP6 overexpression vector (KELD6-2-D6F) (n=1). (C) KELLY, KELD6-2-E and KELLY-2-D6F cells were treated were increasing concentration of lorlatinib and crizotinib and cell viability measured (n=3).

### 3.5 ALK inhibition has greater phosphoproteomic response in DUSP6-depleted cells

A recent study demonstrated DUSP6 is able to target DRP1 independently of ERK [35] and directly dephosphorylate NOTCH [37]. Considering that our data suggests that KELLY cells can dispense with DUSP6 in their control of ERK activity, we hypothesised that DUSP6 should have alternative targets and functions downstream or alongside of ALK. To identify other potential DUSP6 substrates or effector pathways, a phosphoproteomic screen was performed on KELLY cells with and without DUSP6 (Figure 6A). Prior to cell lysis, cells were also treated with lorlatinib or solvent. We used a mixed CRISPR/Cas9-targeted population (KELLY-G+dox) rather than a subclone, as we were aware of subclone heterogeneity (Figure 5). Western blot analysis demonstrated DUSP6 expression was effectively abolished in this mixed population of cells with the *DUSP6* gRNA (Supplementary Figure 6). Volcano plots were generated for each phosphoproteomic comparison, revealing the proportion of significantly upregulated and downregulated phosphopeptides (Figure 6B). Interestingly, loss of DUSP6 increased the number of upregulated and downregulated phosphopeptides in response to ALK inhibition by 4-fold and 8-fold, respectively (Figure 6C). This suggests DUSP6 has a broad influence over ALK signaling in KELLY cells.

**Figure 6.**
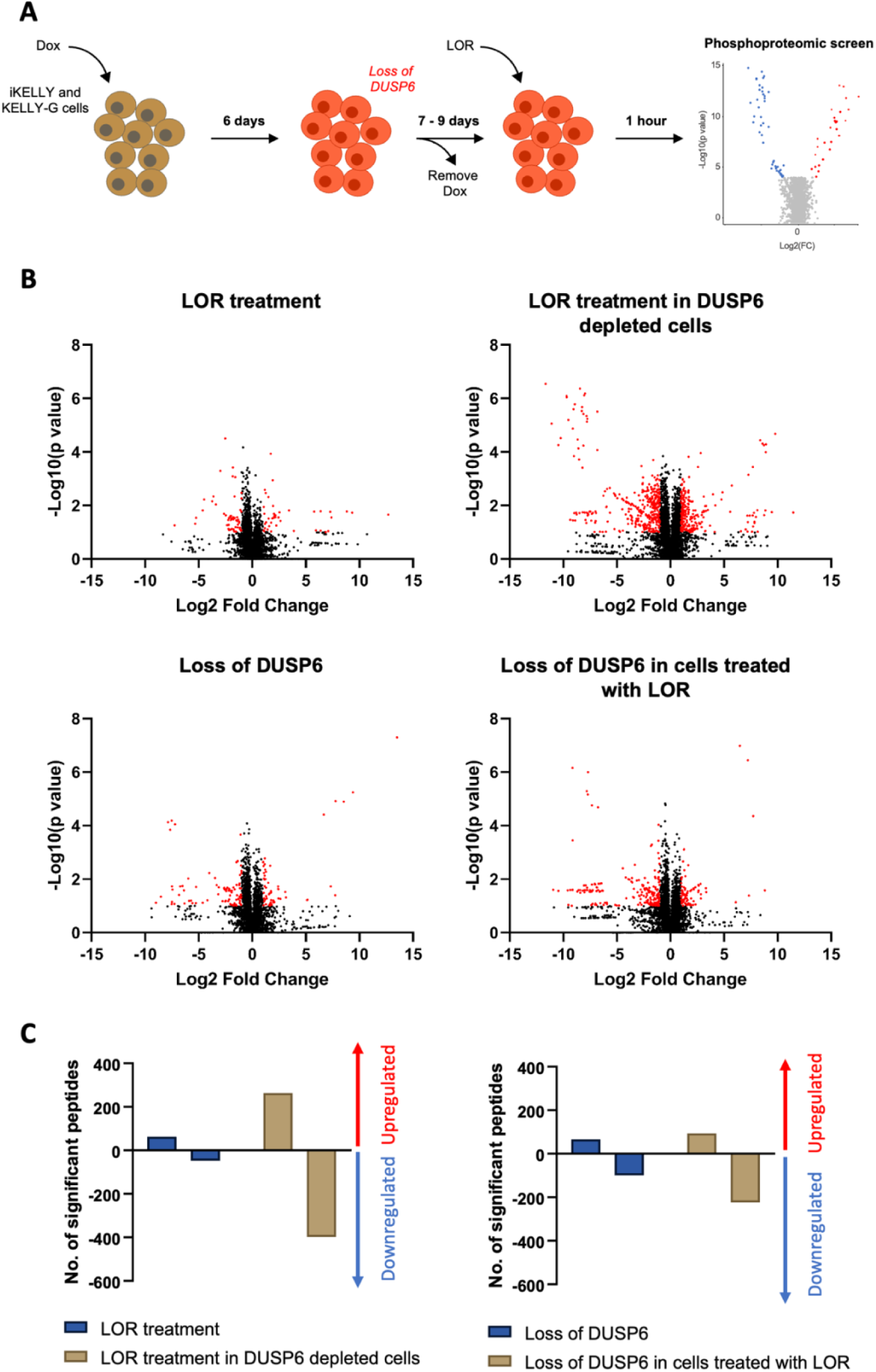
ALK inhibition has a greater effect on the KELLY phosphoproteome in DUSP6-depleted cells. (A) Schematic depicting protocol for phosphoproteomic screen. iKELLY and KELLY-G cells were treated with dox to deplete DUSP6 protein. This cell population was then subsequently treated in replicates with 30 nM lorlatinib for 1 hour and lysates underwent pTyr and pSer/Thr phosphoproteomic analysis (n=4). (B) Volcano plots for four comparison groups where each point represents a phosphosite. Significant thresholds for differential phosphorylation were set as Log2FoldChange > 1 and p-value < 0.1 (red points). (C) The frequency of upregulated and downregulated significant phosphopeptides for each comparison.

To determine the pathways affected by loss of DUSP6, a KSEA analysis was performed (Figure 7; total analysis in repository 10.17632/f2jx8t4x8h.1 [Mendeley Data]). ERK pathway components such as MAP2K1 (MEK) and JAK2 were significantly downregulated by lorlatinib treatment, concordant with the loss of phosphorylation in ERK1/2 peptides (T185, Y187, T190, Y204; see repository 10.17632/f2jx8t4x8h.2), and in accord with previous immunoblots demonstrating lorlatinib-mediated loss of ERK activity (Figure 3B). However, these pathways were relatively insensitive to the removal of DUSP6, perhaps indicating that DUSP6 has alternative targets. Other reported ALK pathway components such as AKT, STAT3 and ERK5 did not appear altered, suggesting ALK signals predominantly via the MEK/ERK pathway in KELLY cells under these conditions [38–40]. The phosphopeptides derived from activated AKT, JNK and p38 were also either undetected or unchanged after ALK inhibition in the *DUSP6*-null subclones (data in repository 10.17632/f2jx8t4x8h.2). Moreover, in the pathways shown in Figure 7, loss of DUSP6 alone did not result in any significant pathway changes that were not also seen with ALK inhibition (for example CSNK2A1, CDK1 and CDK2), indicating a potential level of co-regulation of these. To support this, combining ALK inhibition with DUSP6-deficiency generated little further change in these particular pathways, suggesting that ALK and DUSP6 may act epistatically in the same pathway. There were several pathways that, although weakly affected by ALK inhibition alone or DUSP6 loss alone, were in contrast significantly affected by combined loss of both. These pathways may require additive or synergistic cooperation between ALK and DUSP6. One of these, the EEF2K pathway, was strongly suppressed when both activities were suppressed. Interestingly, loss of EEF2K expression in *MYCN*-amplified neuroblastoma cells exposes them to the cytotoxic effects of nutrient starvation [41]. Overall, these data suggest that DUSP6 can modulate several pathways that are influenced by ALK in this ALK-addicted KELLY line.

**Figure 7.**
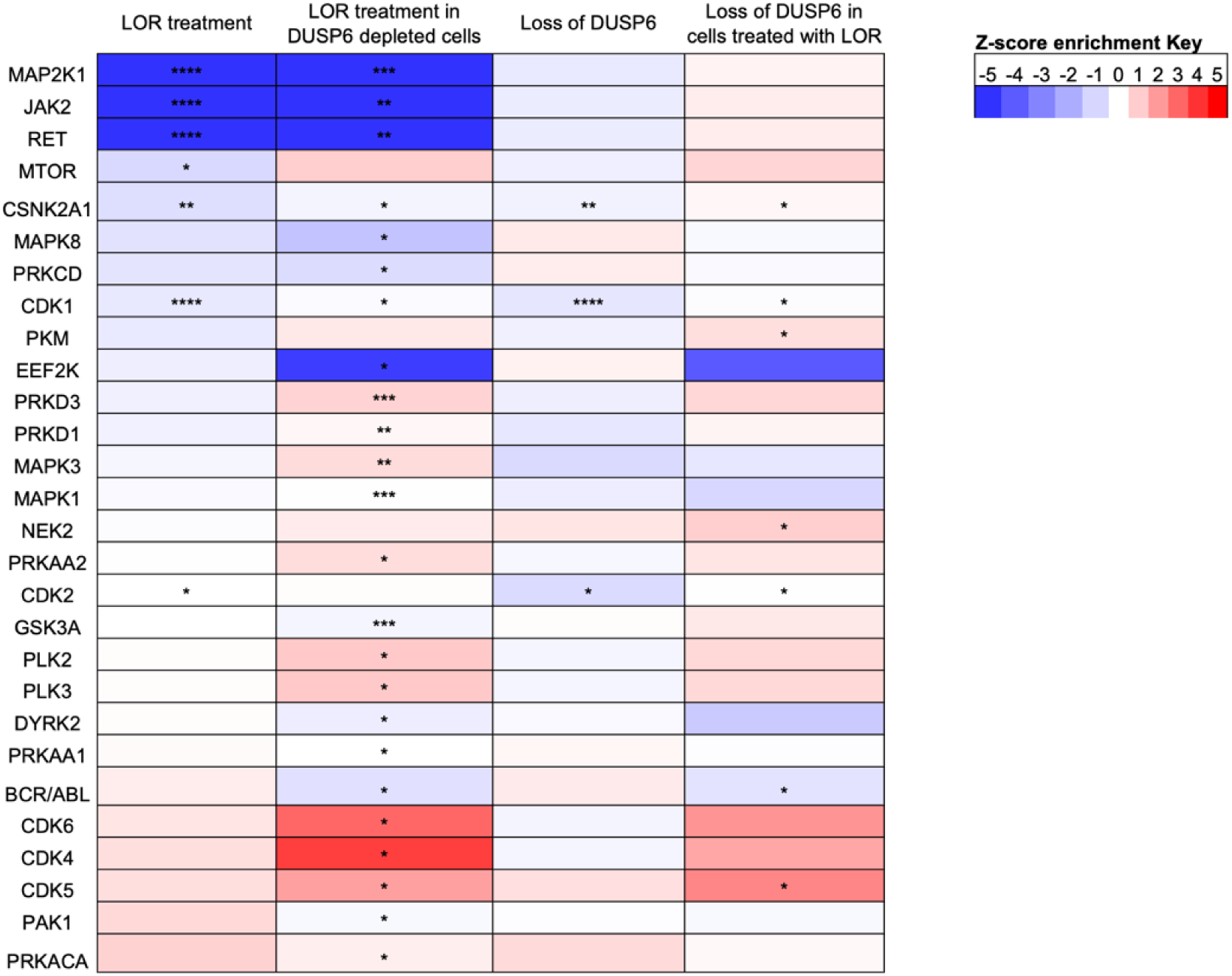
Pathway analysis of phosphoproteomic database after inhibition of ALK. (A) KSEA heatmap of significantly enriched pathways for each treatment comparison [24]. Red and blue indicate potential activation and inactivation of kinases, respectively. Only pathways containing a statistically significant event is included. The p-value represents a statistical assessment of the z-score where *p ≤ 0.05; **p ≤ 0.005; ***p ≤ 0.0005.

### 3.6 DUSP6 may cooperate with ALK to maintain MYCN activity

ALK is known to cooperate with MYCN to drive a subset of aggressive neuroblastoma [42]. N-Myc protein stability is regulated by sequential phosphorylation events on Ser62 and T58, leading to protein degradation, and these are regulated by pathways involving MTOR, PI3K, ERK1/2, GSK3b and CDK1 [43–45]. ALK is considered to increase N-Myc stability through its activation of PI3K, which in turn can inhibit GSK3b and block T58 phosphorylation [46, 47]. In the KSEA data, there are effects on some of these pathways, although the net effect is not clear (Figure 7). Here we sought to understand if DUSP6 functions alongside ALK in regulating N-Myc phosphorylation state and protein level. In the phosphoproteomic screen, one peptide, KFELLPTPPLSPSR, contains both phosphosites T58 and S62, and we assume here given the overexpression of MYCN, that this arises from MYCN not MYC. Three peptides were identified corresponding to single phosphorylated T58 and or dual phosphorylated S62/S64 and T58/S64 (Figure 8A). Sole phosphorylated peptides with either S62 or S64, or dual phosphorylated T58/S62 were not identified. Inhibition of ALK reduced the MYCN dual S62/64 phosphorylated form, with DUSP6 loss having no effect. The role of S64 phosphorylation on MYCN function is unclear. Unexpectedly and in contrast to the work in animal models [46], T58 phosphorylation, which can lead to proteolysis of MYCN, shows potential loss after ALK inhibition (Fig8A). However, our phosphoproteomics data could neither confirm AKT activation nor GSK3b inhibition at their defined phospho-activation sites (data in repository 10.17632/f2jx8t4x8h.1). T58 is increased basally in the cells when DUSP6 activity is absent, whereas absence of DUSP6 reverses the loss of T58 or T58/S64 caused by 1hr inactivation of ALK. It thus appears that DUSP6 does alter the dynamics of MYCN phosphorylation in KELLY cells, somewhat countering ALK in the phosphorylation of these peptides, but it is unclear what the net result on MYCN protein levels would be. We looked at N-Myc protein expression directly in lysates after treatment with 1 hr of lorlatinib. Three of these 4 samples used for proteomics could be analysed in immunoblots. Lorlatinib reduced MYCN expression in cells with depleted DUSP6 in each experimental replicate (Figure 8B). Although not statistically significant, this suggests that although DUSP6-deficiency alone does not alter MYCN levels on average, cells may become sensitised to MYCN loss during concurrent loss of ALK activity. Furthermore, a longer 3 hr treatment with lorlatinib did reduce MYCN protein after ALK inhibition in DUSP6-containing cells, but this occurred to a greater extent in the DUSP6-deficient subclone KELD6-1 (in 3 of 4 replicates) (Figure 8C). These data tentatively suggest again that DUSP6 may operate with ALK to optimally promote or stabilise MYCN expression in some neuroblastoma cells, although the mechanism remains unclear.

**Figure 8.**
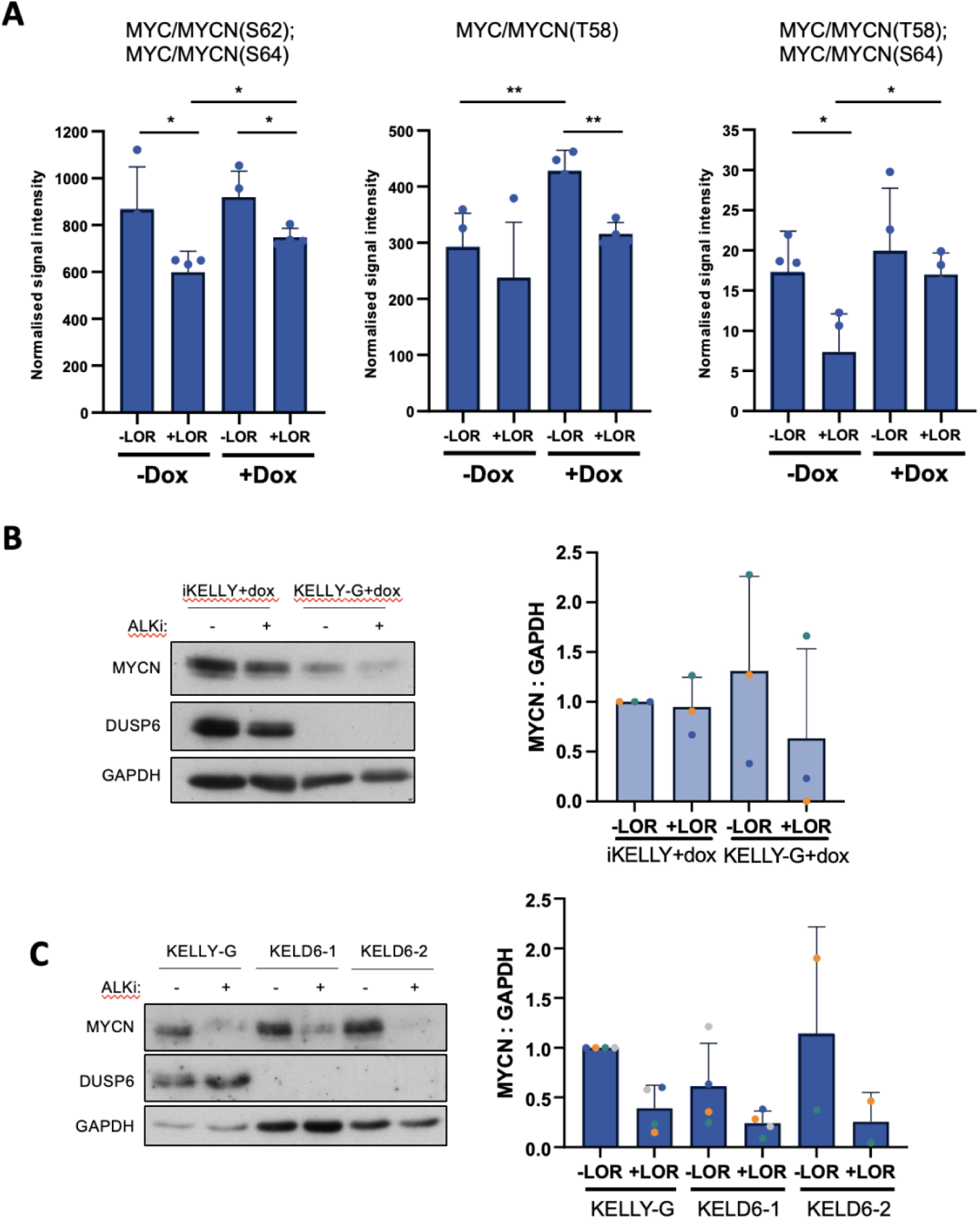
DUSP6 may promote ALK-mediated stabilisation of MYCN. (A) Raw intensity values for three MYC/MYCN phosphopeptides corresponding to Ser62, Ser64 and Thr58. Independent samples t-test compared to 1 hr lorlatinib treatment; *p ≤ 0.05; **p ≤ 0.005. (B) Western blot performed on 3/4 samples sent for phosphoproteomic sequencing to assess total MYCN protein after treatment with 30nM lorlatinib for 1 hour in cells with depleted DUSP6 expression. (C) KELLY-G and *DUSP6*-KO subclones were treated with 30 nM lorlatinib for 3 hours and MYCN expression was assessed by Western blot (n = 2 or 4). KELLY-G was not pretreated with dox therefore DUSP6 is expressed.

## 4. Discussion

Molecular therapies targeting the ALK oncogene display great promise, with the 3^rd^ generation inhibitor lorlatinib currently in clinical trials for cancers including neuroblastoma. Due to potential resistance to single agent ALK therapies, however, it will be of value to understand whether additional enzymes can cooperate or synergise with ALK and thus offer additional therapeutic opportunities. Tyrosine kinases such as ALK are key activators of ERK1/2 pathways and ERK1/2 is in turn usually inactivated by DUSP enzymes through feedback regulation [13, 14]. DUSP6 is a prime example of such a enzyme and it might be expected to be under significant control of the RAS/ERK signal downstream of ALK. DUSP6 function, however, is highly dependent on tumour cell context, showing pro- and anti-oncogenic in different models [15]. By examining the role of DUSP6 in the context of ALK oncogene action in neuroblastoma cells, we have discovered that DUSP6 may in fact act on concert with ALK when ALK is mutated, promoting the latter’s oncogenic potential.

We have found that *DUSP6* mRNA and protein levels are highly dependent on the sustained activity of activated ALK in KELLY, SY5Y and LAN5, but not in non-ALK-mutated IMR32 and SKNAS. Related transcriptional effects have been observed in a range of neuroblastoma lines and with DUSP4 protein in LB-BAR, CLB-G cells [12, 32, 33]. In our study, this expression appears to be largely under the transcriptional rather than post-transcriptional control of ALK in KELLY cells, although LAN5 cells did demonstrate the potential role of ALK in maintaining a proteolytic pathway for DUSP6. MTOR can drive DUSP6 proteolysis independently of ERK in CCL39-derived hamster fibroblast cells, therefore this could be investigated in ALK-mutated neuroblastoma [34]. The levels of DUSP6 also do not straightforwardly correlate with the levels of ERK activation in SY5Y, KELLY and LAN5. In KELLY and LAN5 it appears that ALK controls a second pathway to sustain DUSP6 levels independently of MEK/ERK. In IMR32 and SKNAS, however, DUSP6 expression was more directly correlated with ERK activation level, suggesting that in ALK-mutated cells addicted to this kinase, the normal regulatory circuits between DUSP6 enzymes and ERK activation can be partially bypassed. This novel finding can be extended with data from the *DUSP6* knockout cells, where DUSP6 depletion shows little if any effect on p-ERK levels in KELLY cells or IMR-32 cells even after EGF stimulation. The commonly described feedback regulation between DUSP6 and ERK is therefore not obvious in these cells and might be substituted by another DUSP member.

We observed a modest reduction in cell proliferation in mixed KELLY populations lacking *DUSP6* expression after CRISPR/Cas9 knockdown and subclones showed some variable reductions in proliferation. IMR-32 cells, not addicted to ALK action, shown no such alterations. By itself, therefore, DUSP6 may only influence proliferation weakly and potentially only in cell lines with ALK-activation, although this would require further study to confirm. Nevertheless, loss of DUSP6 in KELLY subclones did sensitise them to ALK inhibitors. This effect may also be an underestimate, given that as ALK inhibition increases, *DUSP6* mRNA and protein levels drop significantly in the wild type cells as well. DUSP6 may thus be able to cooperate with ALK and potentially increase the cell’s addiction to this oncoprotein. This sensitization was only weakly rescued by overexpression of DUSP6, however, so this rescue needs further investigation with more precisely controlled levels of ectopic DUSP6.

Phosphoproteomic profiling allowed a more detailed examination of the biochemical consequences of DUSP6 deficiency in KELLY cells. DUSP6-deficient cells compared to WT cells elicited a very different phosphoproteomic response to ALK inhibition, both quantitatively and qualitatively, indicating that DUSP6 has a significant, albeit complex influence over ALK signaling in these cells. Once again, little evidence was found that DUSP6 influences ERK1/2 activation state, although there was a weak indication of ERK1/2 pathway activation in DUSP6-deficient cells from KSEA analysis. There was also no indication that DUSP6 influenced AKT, p38 or JNK, in agreement with the previous biochemical data. Perhaps surprisingly, there was no evidence of these pathways being maintained by ALK either (data not shown). Instead, ALK inhibition predominantly suppressed MAP2K1, JAK2 and RET kinase pathways from KSEA analysis, based on the MAPK1/3 suppression, and weaker suppression in MTOR, CSNK2A1 and CDK1 pathways. Several pathways showed evidence of either additive or synergistic interactions between ALK and DUSP6, with greater pathway changes when both ALK and DUSP6 were suppressed. One of strongest of these was that of EE2FK signaling, which was reduced in strength only with the double loss of ALK and DUSP6. One phosphosite affected was serine 445, a known autophosphorylation and activating event for EE2FK [48]. EE2FK activation reduces translation in cells, whereas DUSP6 and ALK loss together should logically then increased translation. It is interesting that EE2FK activity has been implicated in promoting the survival of MYC-amplified neuroblastoma cells under nutrient stress conditions, where EE2FK may protect the tumour cells from excessive translation [41]. It remains for future study to understand if EE2FK activity is indeed reduced in cells lacking ALK and DUSP6 activities, and if this triggers increased translation that is detrimental to the cells, or cells become more nutrient-dependent. One further protein of interest may be TWIST1, which promotes invasiveness and poor prognosis in neuroblastoma tumours [49] and cooperates with MYCN to suppress apoptosis [50]. In the phosphoproteomic data, S68 phosphorylation on TWIST1 was suppressed by log1.35 (p<0.01) when DUSP6-deficient cells were treated with lorlatinib, but not in DUSP-expressing cells. Ser68 is a stabilising and activating site and a known target of MAP kinases [51, 52] and so the cooperation of DUSP6 and ALK in potentially promoting Ser68 phosphorylation on TWIST may be of future interest.

One well known oncoprotein that synergises with ALK is N-Myc. *MYCN* amplification drives poor prognosis in children, and co-activation of ALK leads to some of the worst patient outcomes. N-Myc is tightly controlled by proteolysis through sequential phosphorylation on ser62 by CDK1 and other kinases, then thr58 by GSK3β, followed by ser62 dephosphorylation by PP2A and MYCN proteolysis [47]. We did not observe changes in phosphorylation of these kinases and phosphatases after loss of ALK or DUSP6 activities. ALK inhibition did reduce MYCN levels after 3 hr but not at 1hr, with a counterintuitive decrease in Thr58 level after 1hr of treatment. We observed that loss of DUSP6 could stably elevate Thr58 levels, although there was no obvious link to MYCN levels. We did not observe peptides with both ser62 and T58, or ser62 alone, although we did observe double phosphorylated peptides with ser64. Ser64 is a highly conserved site in the MYCbox1 domain of MYCN, but its role is not known. From the overall pattern of N-Myc phosphopeptides therefore, we can say that DUSP6 generally counteracts the effects of ALK inhibition. The net effects of the peptide changes observed are not yet clear, although the doubly DUSP6-deficient and ALK-inhibited cells have on average the least N-Myc protein in them, so the net activities of DUSP6 and ALK could still be cooperating to maintain N-Myc levels.

It remains unclear what the direct targets are for DUSP6 in KELLY cells and how DUSP6 signaling overlaps with that of ALK. Although DUSP6 can be activated by interactions with the ERK kinase interaction motif, there is little clear link between DUSP6 level and ERK1/2 activity in ALK-addicted cells. There is other evidence that DUSP6 can function on other targets [35, 53, 54] [37]. For example, DUSP6 dephosphorylates dynamin-related protein 1 (DRP1 or DNM1L), a protein that regulates mitochondrial fission [35]. In vitro, DUSP6 phosphatase activity was increased by incubation with DRP1, suggesting ERK is not the only protein capable of activating DUSP6. Furthermore, in HeLa cells, overexpression of DUSP6 dephosphorylated Drp1 ser616 regardless of whether ERK was stimulated or inhibited. Curiously, in our proteomics data, we see a small but significant rise in DRP1 ser616 after inhibition of ALK in DUSP6-deficient cells, but not simply through loss of DUSP6, suggesting that DRP1 is one among many of the proteins that is modulated directly or indirectly by this pair of enzymes.

## 5. Conclusion

This study has shown that the DUSP6 enzyme can work alongside the ALK oncoprotein kinase in neuroblastoma cells, acting in several ways as a cooperating enzyme rather than an antagonist, and acting in a pro-oncogenic manner. DUSP6 protein levels are highly dependent on active ALK signaling, but this appears to be at a transcriptional level. DUSP6 also does not appear to be part of the expected ERK1/2 regulatory circuit in KELLY cells at least, but DUSP6 loss leads to broad changes in the cellular phosphoproteome, including the phosphorylation of MYCN, when ALK is co-inhibited. These data suggest that the direct targets of DUSP6 in neuroblastoma cells will be worth identifying and could help us further understand the basis of ALK signaling in these tumour cells.

## Abbreviations

DUSP: dual specificity phosphatase
Dox: doxycycline
CHX: Cycloheximide
LOR: Lorlatinib
CRIZ: Crizotinib
KSEA: Kinase substrate enrichment analysis

## Acknowledgements and Funding

ET was funded by a PhD Studentship from the Child Health Research CIO Charity (Charity number 1152623). AWS and VP were funded by Neuroblastoma UK (Stoker2016NBUK). AWS was funded by the Olivia Hodson Cancer Fund, Great Ormond Street Hospital Children’s Charity (SR16A51). The study was also supported by the NIHR Great Ormond Street Hospital Biomedical Research Centre; the views expressed are those of the author(s) and not necessarily those of the NHS, the NIHR or the Department of Health. Funding for VR and PRC was from Blood Cancer UK (ref. 20008).

## Author contributions

ET: Conceptualization, Data curation, Formal analysis, Investigation, Methodology, Project administration, Validation, Visualization, Writing – original draft & editing.

VP: Conceptualization, Data curation, Formal analysis, Investigation, Methodology, Project administration, Validation.

TK: Data curation, Formal analysis, Investigation, Methodology.

VR and PRC: Data curation, Methodology, Software, Validation.

AWS: Conceptualization, Data curation, Formal analysis, Funding acquisition, Investigation, Methodology, Project administration, Resources, Supervision, Validation, Visualization, Writing – original draft & editing.

## Open Access

For the purpose of open access, the authors have applied a Creative Commons Attribution (CC BY) licence to this preprint and to any Author Accepted Manuscript version arising from this preprint. The phosphoproteomics data that support the findings of this study are openly available in Mendeley Data repository at https://data.mendeley.com/drafts/f2jx8t4x8; DOI 10.17632/f2jx8t4x8h.2. Much of the remaining data in this work is also freely accessible in the PhD thesis of ET published at https://discovery.ucl.ac.uk/id/eprint/10148906.

## References

1. Brodeur, G.M., Neuroblastoma: biological insights into a clinical enigma. Nature Reviews Cancer, 2003. 3(3): p. 203 – 216.

2. Park, J.R., et al., Children’s Oncology Group’s 2013 blueprint for research: neuroblastoma. Pediatr Blood Cancer, 2013. 60(6): p. 985–93.

3. Qiu, B. and K.K. Matthay, Advancing therapy for neuroblastoma. Nat Rev Clin Oncol, 2022. 19(8): p. 515–533.

4. Brodeur, G.M., et al., Amplification of N-myc in Untreated Human Neuroblastomas Correlates with Advanced Disease Stage. Science, 1984. 224(4653): p. 1121–1124.

5. Chen, Y., et al., Molecular regulation and therapeutic targeting of MYCN in neuroblastoma: a comprehensive review. Front Cell Dev Biol, 2025. 13: p. 1683331.

6. Bellini, A., et al., Frequency and Prognostic Impact of ALK Amplifications and Mutations in the European Neuroblastoma Study Group (SIOPEN) High-Risk Neuroblastoma Trial (HR-NBL1). J Clin Oncol, 2021: p. Jco2100086.

7. Brenner, A.K. and M.W. Gunnes, Therapeutic Targeting of the Anaplastic Lymphoma Kinase (ALK) in Neuroblastoma-A Comprehensive Update. Pharmaceutics, 2021. 13(9).

8. Goldsmith, K.C., et al., Lorlatinib with or without chemotherapy in ALK-driven refractory/relapsed neuroblastoma: phase 1 trial results. Nat Med, 2023. 29(5): p. 1092–1102.

9. Trigg, R.M. and S.D. Turner, ALK in Neuroblastoma: Biological and Therapeutic Implications. Cancers (Basel), 2018. 10(4).

10. Berlak, M., et al., Mutations in ALK signaling pathways conferring resistance to ALK inhibitor treatment lead to collateral vulnerabilities in neuroblastoma cells. Mol Cancer, 2022. 21(1): p. 126.

11. Eleveld, T.F., et al., Relapsed neuroblastomas show frequent RAS-MAPK pathway mutations. Nat Genet, 2015. 47(8): p. 864–71.

12. Lambertz, I., et al., Upregulation of MAPK Negative Feedback Regulators and RET in Mutant ALK Neuroblastoma: Implications for Targeted Treatment. Clinical Cancer Research, 2015. 21(14): p. 3327–3339.

13. Shojaee, S., et al., Erk Negative Feedback Control Enables Pre-B Cell Transformation and Represents a Therapeutic Target in Acute Lymphoblastic Leukemia. Cancer Cell, 2015. 28(1): p. 114–28.

14. Ito, T., et al., Paralog knockout profiling identifies DUSP4 and DUSP6 as a digenic dependence in MAPK pathway-driven cancers. Nat Genet, 2021. 53(12): p. 1664–1672.

15. Ahmad, M.K., et al., Dual-specificity phosphatase 6 (DUSP6): a review of its molecular characteristics and clinical relevance in cancer. Cancer biology & medicine, 2018. 15(1): p. 14 – 28.

16. Xiao, W., et al., DUSP6 promotes motility, invasion, and tumorigenicity of thyroid cancer cells via IL8-induced neutrophil extracellular traps. Sci Rep, 2026.

17. Huang, J., et al., A novel prognostic biomarker DUSP6 promote the malignant progression of bladder cancer through mTOR mediated mitophagy. Front Oncol, 2025. 15: p. 1603069.

18. Su, B.H., et al., OCT4-mediated upregulation of DUSP6 promotes metastasis in non-small-cell lung cancer. J Cancer, 2025. 16(14): p. 4172–4186.

19. Ruckert, M.T., et al., DUSP6 is upregulated in metastasis and influences migration and metabolism in pancreatic cancer cells. Sci Rep, 2025. 15(1): p. 33996.

20. Li, X., et al., IGF2BP3/m6A-mediated intron retention isoform of DUSP6 promotes cisplatin resistance in nasopharyngeal carcinoma by inhibiting pyroptosis. BMC Cancer, 2026. 26(1): p. 214.

21. Momeny, M., et al., DUSP6 inhibition overcomes neuregulin/HER3-driven therapy tolerance in HER2+ breast cancer. EMBO Mol Med, 2024. 16(7): p. 1603–1629.

22. Atabay, E.K., et al., Tyrosine phosphatases regulate resistance to ALK inhibitors in ALK+ anaplastic large cell lymphoma. Blood, 2022. 139(5): p. 717–731.

23. Thompson, E.M., et al., The cytotoxic action of BCI is not dependent on its stated DUSP1 or DUSP6 targets in neuroblastoma cells. FEBS Open Bio, 2022. 12(7): p. 1388–1405.

24. Casado, P., et al., Kinase-substrate enrichment analysis provides insights into the heterogeneity of signaling pathway activation in leukemia cells. Sci Signal, 2013. 6(268): p. rs6; 1–13.

25. Hijazi, M., et al., Reconstructing kinase network topologies from phosphoproteomics data reveals cancer-associated rewiring. Nat Biotechnol, 2020. 38(4): p. 493–502.

26. Alcolea, M.P., et al., Phosphoproteomic analysis of leukemia cells under basal and drug-treated conditions identifies markers of kinase pathway activation and mechanisms of resistance. Mol Cell Proteomics, 2012. 11(8): p. 453–66.

27. Montoya, A., et al., Characterization of a TiO(2) enrichment method for label-free quantitative phosphoproteomics. Methods, 2011. 54(4): p. 370–8.

28. Larsen, M.R., et al., Highly selective enrichment of phosphorylated peptides from peptide mixtures using titanium dioxide microcolumns. Mol Cell Proteomics, 2005. 4(7): p. 873–86.

29. Wittig-Blaich, S., et al., Systematic screening of isogenic cancer cells identifies DUSP6 as context-specific synthetic lethal target in melanoma. Oncotarget, 2017. 8(14): p. 23760–23774.

30. Ekerot, M., et al., Negative-feedback regulation of FGF signalling by DUSP6/MKP-3 is driven by ERK1/2 and mediated by Ets factor binding to a conserved site within the DUSP6/MKP-3 gene promoter. The Biochemical journal, 2008. 412(2): p. 287 – 298.

31. Zeliadt, N.A., L.J. Mauro, and E.V. Wattenberg, Reciprocal regulation of extracellular signal regulated kinase 1/2 and mitogen activated protein kinase phosphatase-3. Toxicology and Applied Pharmacology, 2008. 232(3): p. 408–417.

32. Van den Eynden, J., et al., Phosphoproteome and gene expression profiling of ALK inhibition in neuroblastoma cell lines reveals conserved oncogenic pathways. Sci Signal, 2018. 11(557).

33. Borenäs, M., et al., ALK ligand ALKAL2 potentiates MYCN-driven neuroblastoma in the absence of ALK mutation. The EMBO Journal, 2021: p. e105784.

34. Bermudez, O., et al., Post-translational regulation of the ERK phosphatase DUSP6/MKP3 by the mTOR pathway. Oncogene, 2008. 27(26): p. 3685–91.

35. Ma, R., et al., DUSP6 SUMOylation protects cells from oxidative damage via direct regulation of Drp1 dephosphorylation. Sci Adv, 2020. 6(13): p. eaaz0361.

36. Wu, Y.T., et al., TRIM65 Promotes Invasion of Endometrial Stromal Cells by Activating ERK1/2/C-myc Signaling via Ubiquitination of DUSP6. J Clin Endocrinol Metab, 2021. 106(2): p. 526–538.

37. Png, C.W., et al., DUSP6 regulates Notch1 signalling in colorectal cancer. Nat Commun, 2024. 15(1): p. 10087.

38. Sattu, K., et al., Phosphoproteomic analysis of anaplastic lymphoma kinase (ALK) downstream signaling pathways identifies signal transducer and activator of transcription 3 as a functional target of activated ALK in neuroblastoma cells. FEBS J, 2013. 280(21): p. 5269–82.

39. Umapathy, G., et al., The kinase ALK stimulates the kinase ERK5 to promote the expression of the oncogene MYCN in neuroblastoma. Sci Signal, 2014. 7(349): p. ra102.

40. Moore, N.F., et al., Molecular rationale for the use of PI3K/AKT/mTOR pathway inhibitors in combination with crizotinib in ALK-mutated neuroblastoma. Oncotarget, 2014. 5(18): p. 8737–49.

41. Delaidelli, A., et al., MYCN amplified neuroblastoma requires the mRNA translation regulator eEF2 kinase to adapt to nutrient deprivation. Cell Death & Differentiation, 2017. 24(9): p. 1564–1576.

42. Zhu, S., et al., Activated ALK collaborates with MYCN in neuroblastoma pathogenesis. Cancer Cell, 2012. 21(3): p. 362–73.

43. Sears, R., et al., Multiple Ras-dependent phosphorylation pathways regulate Myc protein stability. Genes Dev, 2000. 14(19): p. 2501–14.

44. Vaughan, L., et al., Inhibition of mTOR-kinase destabilizes MYCN and is a potential therapy for MYCN-dependent tumors. Oncotarget, 2016. 7(36): p. 57525–57544.

45. Gustafson, W.C. and W.A. Weiss, Myc proteins as therapeutic targets. Oncogene, 2010. 29(9): p. 1249 – 1259.

46. Berry, T., et al., The ALK(F1174L) mutation potentiates the oncogenic activity of MYCN in neuroblastoma. Cancer Cell, 2012. 22(1): p. 117–30.

47. Farrell, A.S. and R.C. Sears, MYC degradation. Cold Spring Harb Perspect Med, 2014. 4(3).

48. Pyr Dit Ruys, S., et al., Identification of autophosphorylation sites in eukaryotic elongation factor-2 kinase. Biochem J, 2012. 442(3): p. 681–92.

49. Sepporta, M.V., et al., TWIST1 expression is associated with high-risk neuroblastoma and promotes primary and metastatic tumor growth. Commun Biol, 2022. 5(1): p. 42.

50. Valsesia-Wittmann, S., et al., Oncogenic cooperation between H-Twist and N-Myc overrides failsafe programs in cancer cells. Cancer Cell, 2004. 6(6): p. 625–30.

51. Hong, J., et al., Phosphorylation of Serine 68 of Twist1 by MAPKs Stabilizes Twist1 Protein and Promotes Breast Cancer Cell Invasiveness. Cancer Research, 2011. 71(11): p. 3980–3990.

52. Sun, T., et al., The Small C-terminal Domain Phosphatase 1 Inhibits Cancer Cell Migration and Invasion by Dephosphorylating Ser(P)68-Twist1 to Accelerate Twist1 Protein Degradation. J Biol Chem, 2016. 291(22): p. 11518–28.

53. Lee, B., et al., MicroRNA-211 Modulates the DUSP6-ERK5 Signaling Axis to Promote BRAF(V600E)-Driven Melanoma Growth In Vivo and BRAF/MEK Inhibitor Resistance. J Invest Dermatol, 2021. 141(2): p. 385–394.

54. Ndong, C., et al., Mitogen-activated protein kinase (MAPK) phosphatase-3 (MKP-3) displays a p-JNK-MAPK substrate preference in astrocytes in vitro. Neuroscience letters, 2014. 575: p. 13 – 18.

