## Supplementary data figures for "DUSP6 is transcriptionally upregulated by activated ALK and cooperates with ALK signaling to reduce lorlatinib sensitivity in neuroblastoma cells"

Supplementary Figure 1

#### Example flow chart for generation of DUSP6-deficient cells

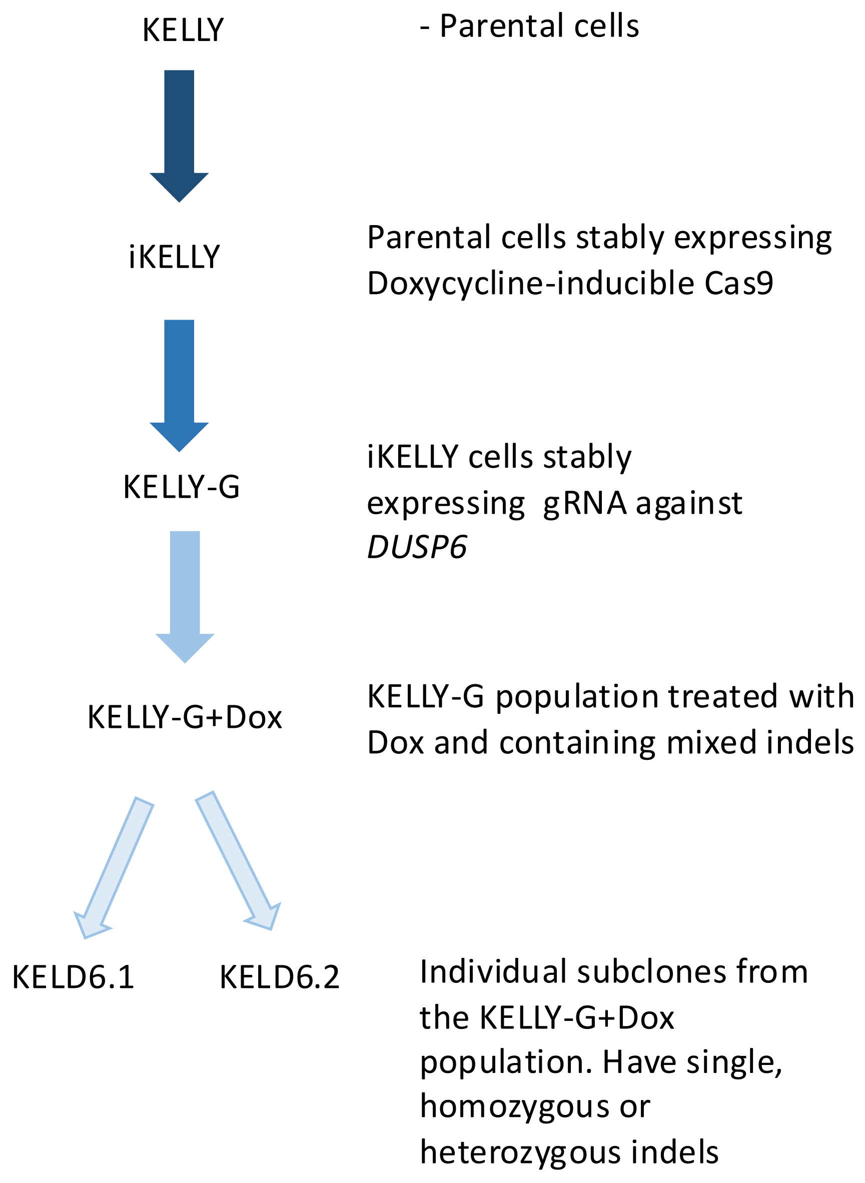

### Supplementary Figure 2

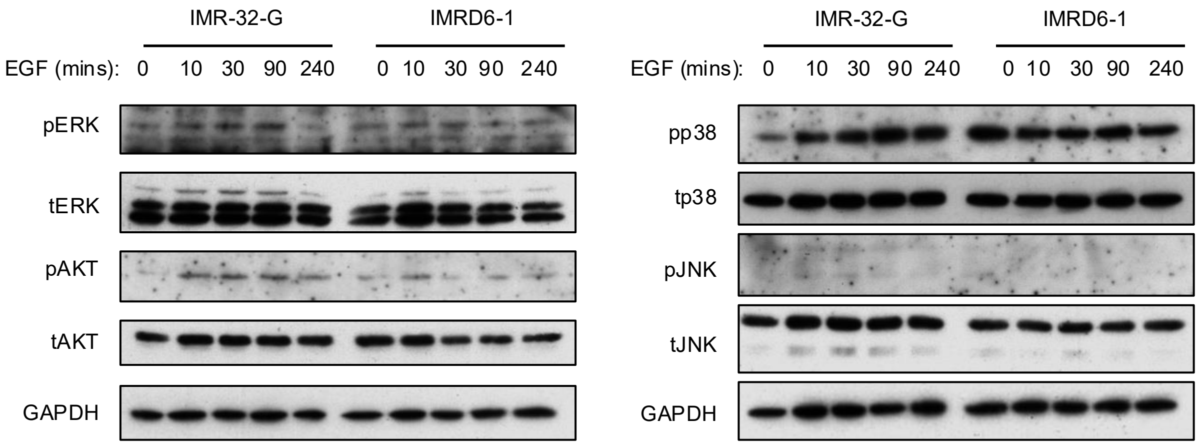


**Supplementary Figure 1 – Loss of DUSP6 has no apparent effect on the MAPK or AKT signalling Pathways in IMR32 cells (preliminary data, n=1)**. IMR32 cells were treated with 30 ng/ml EFG for 10, 30, 90 and 240 minutes and a western blot was performed. IMR-32-G contain the DUSP6 gRNA but are not DOX treated; IMRD6-1 cells are a subclone after DOX treatment, with an indel preventing DUSP6 expression. Although there is no clear change in phosphoprotein levels when DUSP6 is absent, the low phosphoprotein abundance for pERK, pAKT and pJNK prevents accurate quantification.

### Supplementary Figure 3

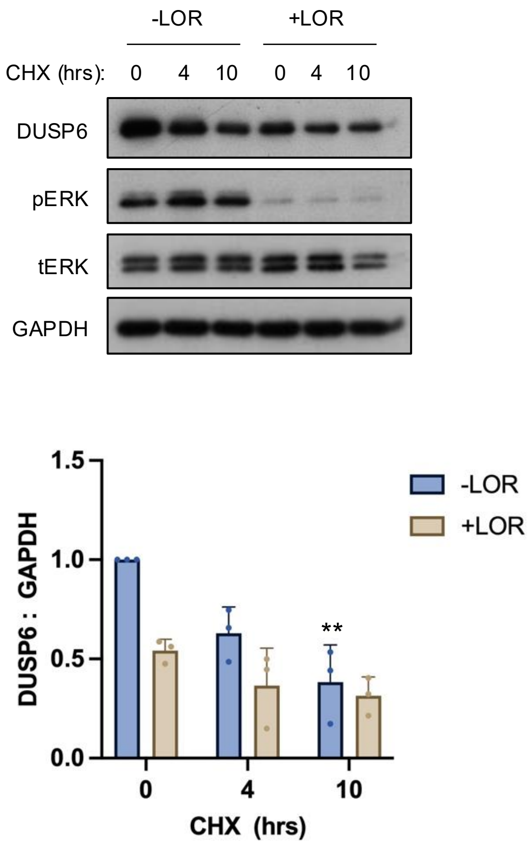


**Supplementary Figure 2. Extended half-life of DUSP6 in KELLY cells.**
KELLY cells were treated with 40 µg/ml CHX for 4 or 10 hrs in the presence or absence of 30 nM Lorlatinib (LOR) followed by immunoblotting for GAPDH, DUSP6, ERK and pERK (n=3). In all case the treatment time with LOR was kept at 10 hours (eg for the 4hr CHX treatment, cells will have received 6hr of prior LOR treatment). For quantification, DUSP6 signal was normalised to GAPDH followed by a second normalisation to the solvent DMSO (-LOR samples). One-way ANOVA compared to DMSO; *$p \leq 0.05$; **$p \leq 0.005$. One-way ANOVA compared to 30 nM LOR treatment; †$p \leq 0.05$; ††$p \leq 0.005$.

### Supplementary Figure 4

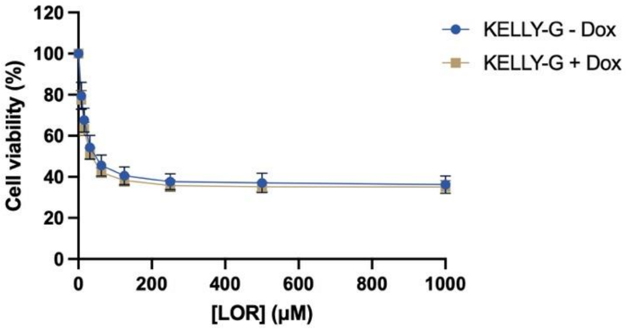


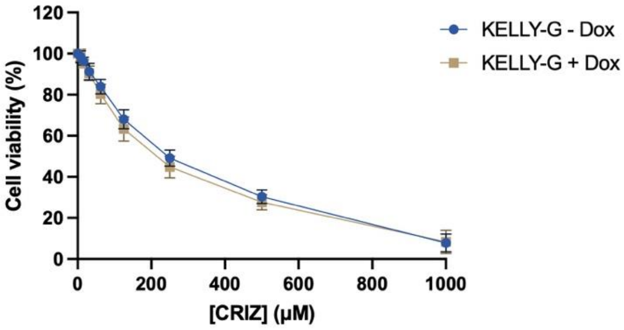


**Supplementary Figure 3. Acute depletion of DUSP6 generates small increase in sensitivity to ALK inhibition.**
KELLY-G cells were incubated with 1 μg/ml Dox (or solvent alone) for 6 days to generate mixed populations with numerous indels. These were then treated with increasing doses of lorlatinib (LOR) and crizotinib (CRIZ) for 7 days. At the end point, cell viability was measure using the resazurin assay (n = 5). Small trends in increased drug sensitivity are seen, but independent samples t-test comparing equivalent drug concentrations did not identify any statistically significant differences.

### Supplementary Figure 5

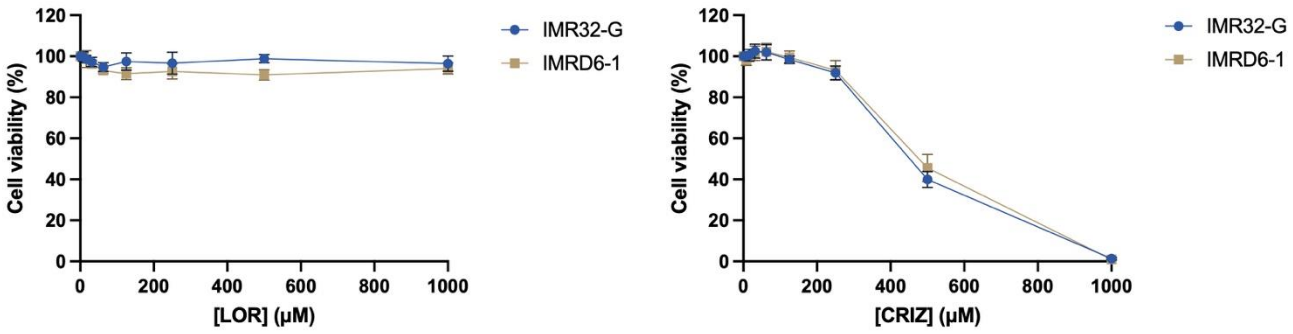


**Supplementary Figure 4. Loss of DUSP6 does not sensitise IMR-32 cells to ALK inhibition.** IMR-32 cells were treated with increasing doses of lorlatinib (LOR) and crizotinib (CRIZ) for 7 days and cell viability was measure using the resazurin assay (n ≥ 3). Independent samples t-test comparing equivalent drug concentrations showed no significant differences. IMR32-G cells contain the DUSP6 gRNA but are not Dox treated; IMRD6-1 are a non-DUSP6 expressing subclone of IMR32-G after Dox treatment and indel formation in *DUSP6*.

### Supplementary Figure 6

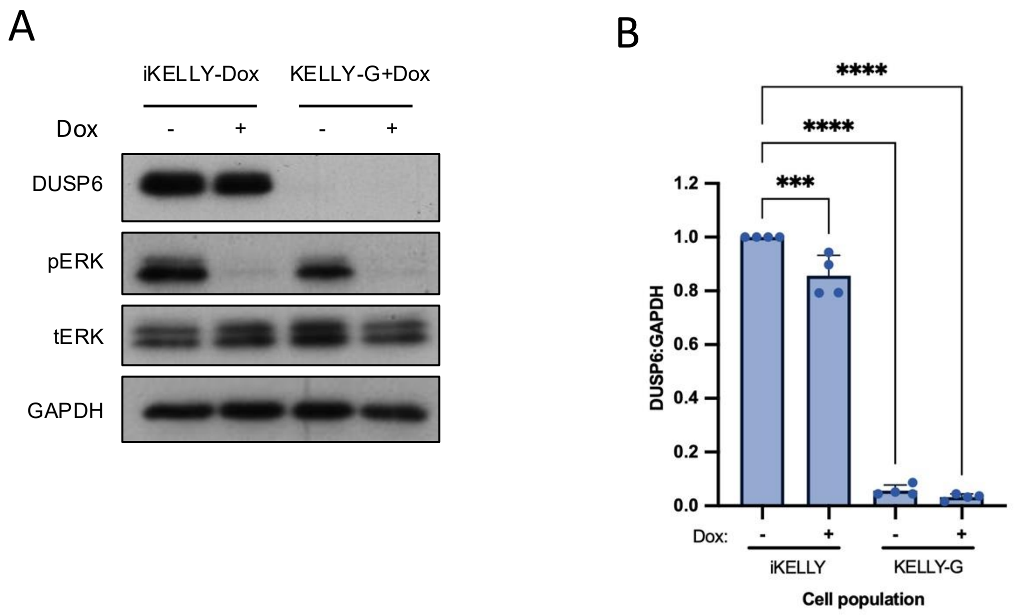


**Supplementary Figure 5. Cell line inputs into phosphoproteomic analysis.**
iKELLY cells (containing inducible Cas9) and KELLY-G (iKELLY containing inducible Cas9 plus the *DUSP6* gRNA) were treated with either DMSO or Dox for 6 days. During this time, Dox induces Cas9 to introduce heterogeneous indels in KELLY-G. Dox was then removed for 7-9 days and cells then treated with lorlatinib (LOR) or solvent for 1 hour before lysing. A phosphoproteomic screen was performed on lysates. **A**, immunoblot for lysates sent for phosphoproteomic analysis, probed for DUSP6, pERK, total ERK and GAPDH. For quantification in **B**, DUSP6 and was normalised to GAPDH, followed by a second normalisation to the iKELLY-LOR sample. **B**, one-way ANOVA: **p $\leq$ 0.005; ***p $\leq$ 0.0005.
